# *Mycobacterium abscessus* biofilms produce an ECM and have a distinct mycolic acid profile

**DOI:** 10.1101/2021.03.05.434154

**Authors:** Anja Dokic, Eliza Peterson, Mario L Arrieta-Ortiz, Min Pan, Alessandro Di Maio, Nitin Baliga, Apoorva Bhatt

## Abstract

A non-tuberculous mycobacterium (NTM), *Mycobacterium abscessus* is an emerging opportunistic pathogen associated with difficult to treat pulmonary infections, particularly in patients suffering from cystic fibrosis. It is capable of forming biofilms *in vitro* that result in an increase of already high levels of antibiotic resistance in this bacterium. Evidence that *M. abscessus* forms biofilm-like microcolonies in patient lungs and on medical devices further implicated the need to investigate this biofilm in detail. Therefore, in this study we characterized the *M. abscessus* pellicular biofilm, formed on a liquid-air interface, by studying its molecular composition, and its transcriptional profile in comparison to planktonic cells. Using scanning electron micrographs and fluorescence microscopy, we showed that *M. abscessus* biofilms produce an extracellular matrix composed of lipids, proteins, carbohydrates and eDNA. Transcriptomic analysis of biofilms revealed an upregulation of pathways involved in the glyoxylate shunt, redox metabolism and mycolic acid biosynthesis. Genes involved in elongation and desaturation of mycolic acids were highly upregulated in biofilms and, mirroring those findings, biochemical analysis of mycolates revealed molecular changes and an increase in mycolic acid chain length. Together these results give us an insight into the complex structure of *M. abscessus* biofilms, the understanding of which may be adapted for clinical use in treatment of biofilm infections, including strategies for dispersing the ECM, allowing antibiotics to gain access to bacteria within the biofilm.

## 1 Introduction

Non-tuberculous mycobacteria (NTM) belong to the genus *Mycobacteria* species and are environmental bacteria capable of causing disease in immunocompromised individuals. NTMs cause skin and soft tissue infections, lymphadenitis and aseptic meningitis, however most commonly they manifest as pulmonary infections (Faria et al., 2015; Ratnatunga et al., 2020). Since the discovery of the association of *Mycobacterium avium* infections with mortality in HIV patients, disseminated and mixed culture NTM infections were found in other immune-compromised populations suffering from chronic pulmonary disease, renal failure, cystic fibrosis (CF), leukaemia and transplant recipients (Sousa et al., 2015).

*M. abscessus* is a rapidly growing NTM and an emerging opportunistic pathogen most often causing chronic pulmonary disease in patients with underlying lung conditions (Medjahed et al., 2010). CF patients are particularly vulnerable, and it is estimated that 3-10% of CF patients in the USA and Europe are infected with *M. abscessus*, which can either result in a poor clinical outcome or life-long persistent infection without symptoms (Cystic Fibrosis Foundation, 2010). *M. abscessus* is the most commonly isolated RGM from lung infections, which is alarming given the average rate of treatment success is only 45.6% (Kim et al., 2016; Kwak et al., 2019). None of the currently available treatments are curative or have been shown as effective in long-term sputum conversion in patients with chronic lung disease (Kwak et al., 2019).

A key strategy of NTM pathogenesis is the ability to form biofilms in the environment as well as in water distribution systems and on hospital equipment (Galassi et al., 2003; Ghosh et al., 2017; Zambrano and Kolter, 2005). Notably, *Mycobacterium fortuitum, Mycobacterium chelonae* and *M. abscessus* subsp. *massilinese* have all been associated with biofilm formation on medical implants (Bosio et al., 2012; Padilla et al., 2018; Rüegg et al., 2015). Biofilms are communities of aggregated bacteria that form in response to stress and as a part of this survival strategy bacteria within the biofilm undergo genetic and metabolic changes (Chakraborty and Kumar, 2019; Solokhina et al., 2017; Vasudevan, 2014). In the host, biofilms contribute to virulence by mediating colonization, evading the host’s immune response and limiting diffusion of antimicrobials (Aung et al., 2017; Faria et al., 2015; Roux et al., 2016).

Bacteria in biofilms are enveloped in an extracellular matrix (ECM) which is required for cell aggregation, adhesion to surfaces, retention of water and nutrient provision. The ECM also serves as physical barrier, allowing bacteria a significant degree of resistance to water treatment, disinfectants and antibiotics (Aung et al., 2017, 2016; Flemming and Wingender, 2010; Galassi et al., 2003; Greendyke and Byrd, 2008). Secreted components that constitute an ECM can be lipids, carbohydrates, proteins or extracellular DNA (eDNA) (Chakraborty and Kumar, 2019; Flemming and Wingender, 2010; Flemming et al., 2016; Karygianni et al., 2020; Vega-Dominguez et al., 2020). In mycobacterial biofilms, lipids, particularly mycolyl-diacylglycerol, mycolic acids (MA) and glycopeptidolipids (Chen et al., 2006; Ojha et al., 2005a; Recht and Kolter, 2001; Yang et al., 2017) are important for biofilm formation and, additionally, strains defective in exporting cell wall lipids have a defect in biofilm formation (Sophia A. Pacheco et al., 2013). Biofilms of *M. tuberculosis* and *M. avium* have been demonstrated to form an ECM in biofilms (Rose et al., 2015; Trivedi et al., 2016), and we have recently shown the same for *M. chelonae* (Vega-Dominguez et al., 2020). Earlier study found that carbohydrates are the major component of the extracellular material of *M. avium, M. gastri* and *M. kansasii*, while in *M. smegmatis* and *M. phlei* carbohydrates are less abundant in comparison to proteins. (Lemassu et al., 1996b). Carbohydrates are major components of *M. tuberculosis* and *M. chelonae* biofilm ECMs (Lemassu et al., 1996a; Trivedi et al., 2016; Vega-Dominguez et al., 2020) and while they seem to play a structural role, the role of proteins has not been fully characterized but there is evidence that treatment with Proteinase K reduces overall biofilm biomass in *M. tuberculosis* (Trivedi et al., 2016). eDNA, identified in biofilms of *M. tuberculosis, M. avium, M. abscessus, M. fortuitum* and *M. chelonae* (Aung et al., 2017; Rose et al., 2015; Rose and Bermudez, 2016; Toyofuku et al., 2016; Trivedi et al., 2016; Vega-Dominguez et al., 2020; Whitchurch et al., 2002), plays a fundamental structural role by promoting attachment to surfaces and facilitating bacterial aggregation (Ibáñez de Aldecoa et al., 2017; Pakkulnan et al., 2019; Rose et al., 2015) as biofilms treated with DNase become less structurally secure and are more vulnerable to antibiotics (Cavaliere et al., 2014; Rose et al., 2015; Tetz et al., 2009; Trivedi et al., 2016).

Biofilms are suspected to play an important role in *M. abscessus* infections and there is evidence of biofilm-like microcolonies formation in patient lungs (Fennelly et al., 2016; Qvist et al., 2015, 2013). It has been demonstrated that *M. abscessus* has a decreased susceptibility to several first-line antibiotics, such as amikacin, cefoxitin, and clarithromycin, when grown in a biofilm *in vitro* (Greendyke and Byrd, 2008; Marrakchi et al., 2014). Previous research suggests that *M. abscessus* smooth colony variants colonize abnormal lung airways in a biofilm, and the spontaneous loss of surface glycopeptidolipids leads to a morphological change to the invasive rough colony type, causing inflammation and invasive disease (Howard et al., 2006; Rhoades et al., 2009).

Studies in *M. avium, M tuberculosis, M. smegmatis* and *M. chelonae* showed that mycobacterial biofilms are rich in carbohydrates, lipids, proteins and eDNA and have an altered transcription profile (Chakraborty and Kumar, 2019; Rose et al., 2015; Trivedi et al., 2016; Vega-Dominguez et al., 2020), however the same has not yet been shown in *M. abscessus*. In this study we focused on understanding biofilm formation in *M. abscessus*, using a pellicular biofilm formed on a liquid-air interface as a model. Following a similar approach used for *M. chelonae* (Vega-Dominguez et al., 2020), utilising confocal laser scanning microscopy, scanning electron microscopy, transcriptomic and biochemical analysis, we have defined the structure, content and expression profiles of *M. abscessus* biofilms. A greater understanding of molecular events that drive biofilm formation and biochemical changes that occur in the cells is needed in order to improve the current strategies for combating *M. abscessus* infections.

## 2 Materials and methods

### 2.1 Culture conditions

*M. abscessus* ATCC 19977 Type strain was used for all experiments. Planktonic bacteria were grown in Sauton’s minimal media supplemented with 0.05% Tyloxapol shaking (180 rpm) at 37°C. For transcriptomic analysis 1 L of culture was grown per timepoint, planktonic t1 was harvested 24 h post inoculation and planktonic t2 at 48 h. For biofilm formation, *M. abscessus* colonies grown on 7H11/OADC were resuspended in sterile 1xPBS and diluted to OD_600_ 0.05 in Sauton’s minimal media and grown stationary at 37°C in a 24-well plate (for lipid analysis and microscopy) or in 75 cm^2^ culture flasks (for transcription analysis). Biofilm cultures were harvested after 5 days (t1) or 7 days (t2).

### 2.2 Scanning electron microscopy

*M. abscessus* biofilms were grown as described above. Using 70% ethanol-sterilized microscopy cover slips and tweezers, the pellicles was collected from the wells and transferred to a new 24-well plate where they were fixed overnight at 4 °C with freshly prepared 2% glutaraldehyde in PBS. Afterwards, samples were prepared using two methods: airdrying or gradient alcohol dehydration. Airdrying: samples were left to airdry on an absorbent tissue. Gradient alcohol dehydration: samples were dehydrated thought a gradient of ethanol solutions (25%, 50%, 75% and 100%) and then airdried. All samples were mounted on SEM stubs, coated with gold and examined in Philips XL-30 (LaB6) with Link Isis EDS.

### 2.3 Confocal microscopy

*M. abscessus* electrocompetent cells (grown to OD_600_ 0.8) were transformed with pMV306_eGFP plasmid containing a zeocin resistance cassette (2.5 kV, 25 mF, 1000 ohms, 1 mm cuvettes) and selected on 25 µg/ml zeocin. *M. abscessus* eGFP strain was used for all florescence microscopy experiments. Pellicles were fixed with 4% paraformaldehyde for 30 min. Samples were stained with Nile Red (1 µM, 30 min), Propidium iodide (15 µM, 30 min), AlexaFlour647 conjugated to Concanavalin A (100 µg/ml, 45 min) and FilmTracer™ SYPRO™ Ruby Biofilm Matrix Stain (10 µg/ml, 45 min). Cells were visualized on TIRF Nikon A1R system with a Ti microscope frame and a 100x/1.4 PlanApo objective. For co-localization analysis and calculation of biovolume, five images selected at random were taken from five distinct biological samples.

### 2.4 Image processing

The data was analyzed using the Icy platform (http://icy.bioimageanalysis.org) (De Chaumont et al., 2012). Protocols plug-in was used to create automated pipelines for each of the analysis, using a similar method as Vega-Dominguez *et al*., 2020 and Pike *et al*., 2017. Images were first denoised using a Gaussian filter and then the threshold was calculated using the Li method through the Thresholder plug-in. Images were binarized to create a region of interest (ROI) from which a signal was extracted for co-localization analysis using the Co-localization Studio plug-in (Publication ID: ICY-H9X6X2) (Lagache et al., 2015). The outputs were Pearson’s and Mander’s coefficient. The ROI Statistics plug-in was additionally used to determine the biovolume (Publication Id: ICY-W5T6J4).

### 2.5 RNA extraction

Total RNA was extracted from the cultures from five different experiments. 600 µl of freshly prepare lysozyme (5 mg/mL in TE pH8.0) was added to ∼200 µl of pellet to digest the bacterial cell wall. Mixture was transferred to a bead beating tube and incubated at 37°C for 30 min. Subsequently 1/10^th^ of 10% SDS, 3M NaCO_3_ (pH 5.2) and 720 µl Acid Phenol (pH 4.2) were sequentially added with 2 min bead beating steps in-between and finally inverting for 1 min at the end. Samples were incubated at 65°C for 5 min and mixed by inverting every 30 sec. Following a 5 min centrifugation at 14000 rpm, aqueous phase was removed to a fresh tube, topped with 600 µl of acid Phenol (pH 4.2) and mixed by inverting. The centrifugation step was repeated, and the aqueous phase topped with 500 µl of chloroform: isoamyl alcohol. The same centrifugation step was repeated and 1/10th volume of NaCO_3_, 3 volumes of 100% ethanol and 1 µl of glycogen was add, mixed well and left to precipitate overnight at −20°C. The precipitate was washed once with 70% ethanol, airdried and resuspended in 30 µl of RNase-free H_2_O. Concentration of RNA was checked on Thermo Scientific™ *NanoDrop*™ and the quality with Agilent 2100 BioAnalyzer™ System using the appropriate cassette (Agilent RNA 6000 Nano Kit). 10 µg of isolated RNA was treated with DNase for 30 min at 37°C and ethanol precipitated overnight at −20°C. Precipitate was resuspended in 30 µl of water, and the concentrate and quality checked as before. For removal of ribosomal RNA, the Ribo-Zero rRNA removal kit (Illumina) was used without deviations from the manufacturer’s instructions. The quality of RNA was checked on a BioAnalyzer Nano chip (Agilent). TruSeq Stranded mRNA kit (Illumina) was used for subsequent steps of cDNA strand synthesis, adenylation of 3’ ends, ligation and PCR amplification. Final dsDNA product was measured on Qubit and based on this concentration samples were prepared for sequencing.

### 2.6 RNA-seq analysis and differentially expressed genes

Samples were pooled and sequenced using an Illumina NextSeq Instrument. Paired-end 75 bp reads were checked for technical artifacts using Illumina default quality filtering steps. Raw FASTQ read data were processed using the R package DuffyNGS as described previously (Vignali et al., 2011). Briefly, raw reads were passed through an alignment pipeline that first filtered out unwanted rRNA transcripts and then the main genomic alignment stage against the genome. Reads were aligned to *M. abscessus* (ASM6918) with Bowtie2 (Langmead and Salzberg, 2012), using the command line option “very-sensitive.” BAM files from the genomic alignment were combined into read depth wiggle tracks that recorded both uniquely mapped and multiply mapped reads to each of the forward and reverse strands of the genome(s) at single-nucleotide resolution. Gene transcript abundance was then measured by summing total reads landing inside annotated gene boundaries, expressed as both RPKM and raw read counts. RNA-seq data (raw fastq files and read counts) have been deposited in the GEO repository under accession number GSE165352.

A panel of 5 DE tools was used to identify gene expression changes between 5-day old biofilms (Biofilm T1) samples and planktonic samples (24 h, t1) or 7-day old biofilms (Biofilm T2) samples and planktonic samples (24 h, t1). The tools included (i) RoundRobin (in-house); (ii) RankProduct (Breitling et al., 2004); (iii) significance analysis of microarrays (SAM) (Tusher et al., 2001); (iv) EdgeR (Robinson and Smyth, 2008); and (v) DESeq2 (Love et al., 2014). Each DE tool was called with appropriate default parameters and operated on the same set of transcription results, using RPKM abundance units for RoundRobin, RankProduct, and SAM and raw read count abundance units for DESeq2 and EdgeR. All 5 DE results were then synthesized, by combining gene DE rank positions across all 5 DE tools. Specifically, a gene’s rank position in all 5 results was averaged, using a generalized mean to the 1/2 power, to yield the gene’s final net rank position. Each DE tool’s explicit measurements of differential expression (fold change) and significance (*P*-value) were similarly combined via appropriate averaging (arithmetic and geometric mean, respectively). Genes with averaged absolute log2 fold change bigger than two and multiple hypothesis adjusted *P*-value < 0.01 were considered differentially expressed.

### 2.7 Metabolic pathway enrichment analysis

We mapped the significantly differentially expressed genes at biofilm t1 and t2 against the most recent genome-scale metabolic network construction of *M. tuberculosis* H37Rv iEK1011 (Kavvas et al., 2018) by identifying orthologs using OrthoVenn2 (Ling Xu et al., 2019 NAR). We used the subsystem definitions outlined in iEK1011 to explore pathway usage at the network level. We identified metabolic pathways that were significantly enriched in the *M. abscessus* biofilm stages (Benjamini Hochberg corrected hypergeometric *P*-value <= 0.05). For these pathways, we calculated the average fold-change of all genes. Additionally, we performed functional enrichment analysis of *M. abscessus* DEG sets with DAVID (Huang et al., 2008 Nat. Protoc.). Only functional terms with adjusted *P*-value<0.05 were considered as overrepresented.

### 2.8 Lipid analysis

Planktonic and biofilm cultures of *M. abscessus*, grown as previously described, were labelled with ^14^C acetic acid (1 µCi/µl PekinElmer) in a ratio of 1 µl:1 ml of culture. Planktonic cultures were labelled at OD 0.8 and harvested at OD 1.2. Pellicles were labelled after 5 days, at early maturation, and harvested after 7 days, at late maturation. Cultures were harvested by centrifugation (4000 rpm, 10 min) in 10 ml glass tubes sealed with polytetrafluoroethane (Teflon®)-lined screw cap, washed with PBS and then dried at 55°C under air flow. Apolar and polar lipid fractions were isolated as described in Besra, 1998 (Besra, 1998). Radioactive counts of ^14^C labelled samples were measured using a scintillation counter and equalized so that 25,000 counts were loaded on each TLC plate (silica, 6.7×6.7 cm). After being run in various solvent systems in order to separate different fractions of the isolated lipids, radioactive TLCs were exposed to Carestream BIOMAX MR film for 48 h. Lipids were separated thrice in direction I in petroleum ether (60-80):acetone (98:2, v/v) and once in direction II in toluene:acetone (95:5, v/v). FAMEs and MAMEs were isolated as described by Besra, 1998 (Besra, 1998). One direction TLC was run in petroleum ether:acetone (19:1, v/v). Subclasses of MAMEs were separated using 2D argentation TLC as described by Singh *et al*. 2016. 75% of the plate was dipped in 10% aqueous solution of AgNO3 and plates were dried at 90°C for 15 mins; samples were resolved in hexane:ethyl acetate (19:1) twice in dimension I and then in petroleum ether:diethyl ether (17:3) three times in dimension II.

### 2.9 Mass spectrometry

After initial characterization using ^14^C-labelled samples, cultures were up-scaled to 3 l for planktonic and 500 ml for pellicles and grown without ^14^C labelling for lipid extraction and analysis by mass spectrometry. A silica column was used to separate FAMEs and MAMEs using a percentage gradient of toluene:ethyl acetate. Relevant fractions were analysed by MALDI-TOF/MS using the Voyager DE-STR MALDI-TOF instrument.

## 3 Results

### 3.1 *M. abscessus* forms an organized biofilm enveloped in a potential ECM

*M. abscessus* can grow as a pellicular biofilm on a liquid-air interface, that is opaque and wrinkly in appearance (**Fig. 1A**). Using scanning electron microscopy, we studied the formation of *M. abscessus* pellicles after a 6-day incubation. Air-dried biofilms revealed a thick substance, likely an ECM, coating the bacteria under which individual cells are not distinguishable (**Fig. 1B and 1D**). When subjected to gradual alcohol dehydration, this layer was stripped from the biofilm revealing single, stacked bacilli arranged in an organized manner allowing for formation of pores and channels, as indicated by white arrows (**Fig. 1C and 1E**).

**Fig. 1.**
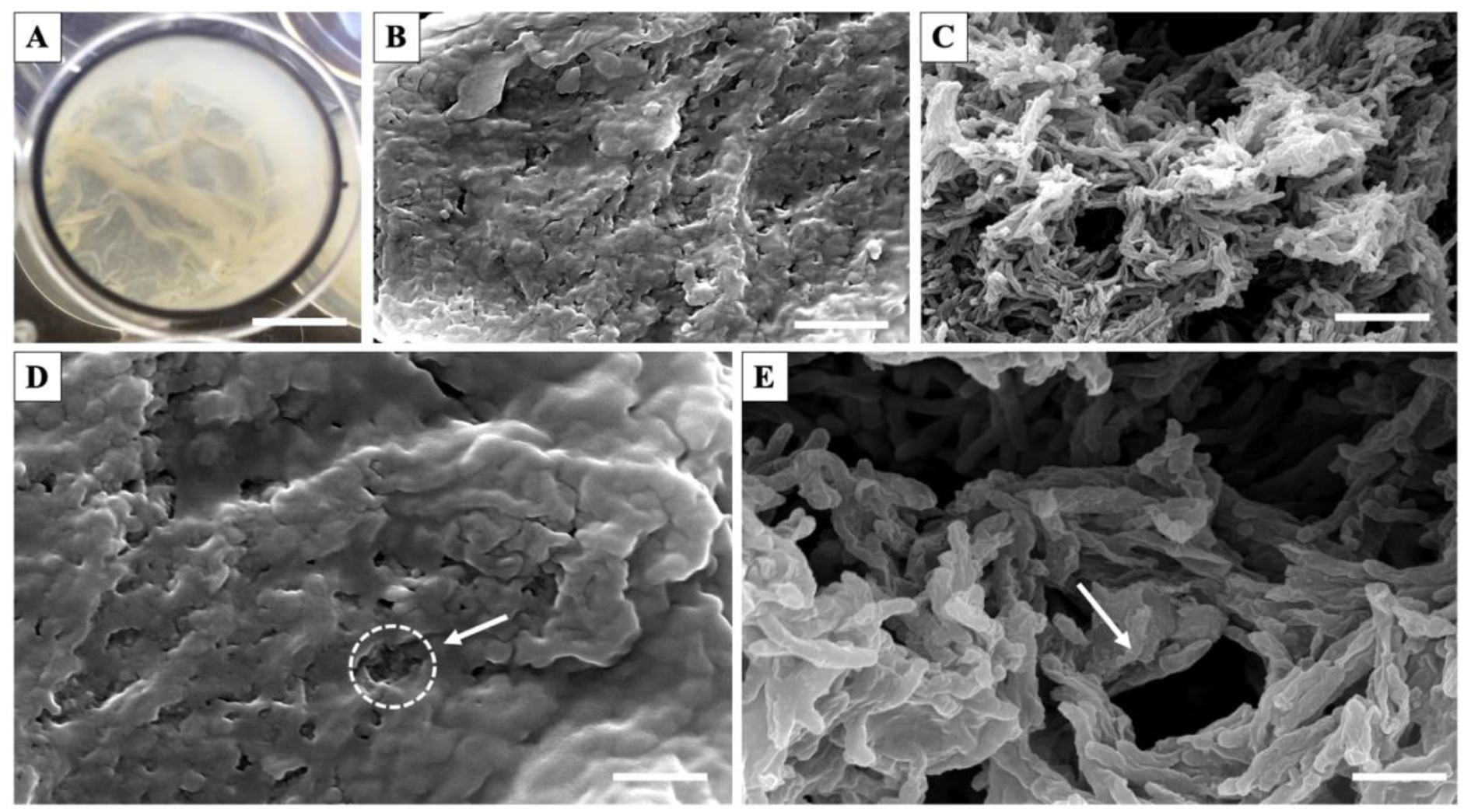
Scanning electron micrographs of *M. abscessus* pellicles grown in Sauton’s media. **A**. 6-day old pellicle in a 24-well plate (*Ø* 15.6 mm, scale bar = 8 mm); **B.-E**. micrographs of *M. abscessus* pellicles under 5000x magnification (**B**. and **C**., scale bar = 5 µm) and 10000x magnification (D. and E., scale bar = 2 µm). Pellicles were airdried (B. and D.) or treated with gradual alcohol dehydration (C. and E.). Circle and arrows point to pores in the ECM.

### 3.2 Fluorescent images revealed the macromolecular content of *M. abscessus* biofilms

Scanning electron microscopy (SEM) studies on the ECM were complemented with confocal laser scanning microscopy (CLSM) using an enhanced green fluorescence protein (eGFP) expressing *M. abscessus* strain and a set of fluorophores to selectively label components of the ECM. Our previous studies with *M. chelonae* biofilms showed that these fluorophores can reveal biofilm matrix composition (Vega-Dominguez et al., 2020). Concanavalin A conjugated with Alexa Flour 647 (AF) was used for staining carbohydrates, Propidium iodide (PI) for nucleic acids, FilmTracer™ SYPRO® Ruby (SR) for proteins and Nile red (NR) for lipids.

Confocal microscopy revealed the presence of proteins, eDNA, lipids and carbohydrates in the *M. abscessus* biofilm matrix (**Fig. 2**). A visual assessment of the micrographs indicated that lipids were the most abundant biomolecule in the biofilms, while proteins, eDNA, and carbohydrates were present, but more dispersed throughout the biofilm structure. Fluorescent images were quantified by calculating the relative volume of each component and evaluating co-localization of the emitted signals of GFP and each dye using Pearson’s and co-occurrence using Mander’s coefficients (Schlafer and Meyer, 2017). Pearson’s correlation is a well-established method for presenting correlation and has been adapted to measure co-localization between fluorophores by showing the degree to which the signals linearly correlate with each other. A value of 1 (+/-) indicates a perfect positive or negative relationship, while 0 shows absence of a relationship. A high Pearson’s correlation coefficient (PCC), denoted with *r*, indicates that both signals from GFP and a fluorophore increase or decrease proportionally, while a low coefficient denotes that signal intensity of one does not alter with the change in intensity of the other.

**Fig. 2.**
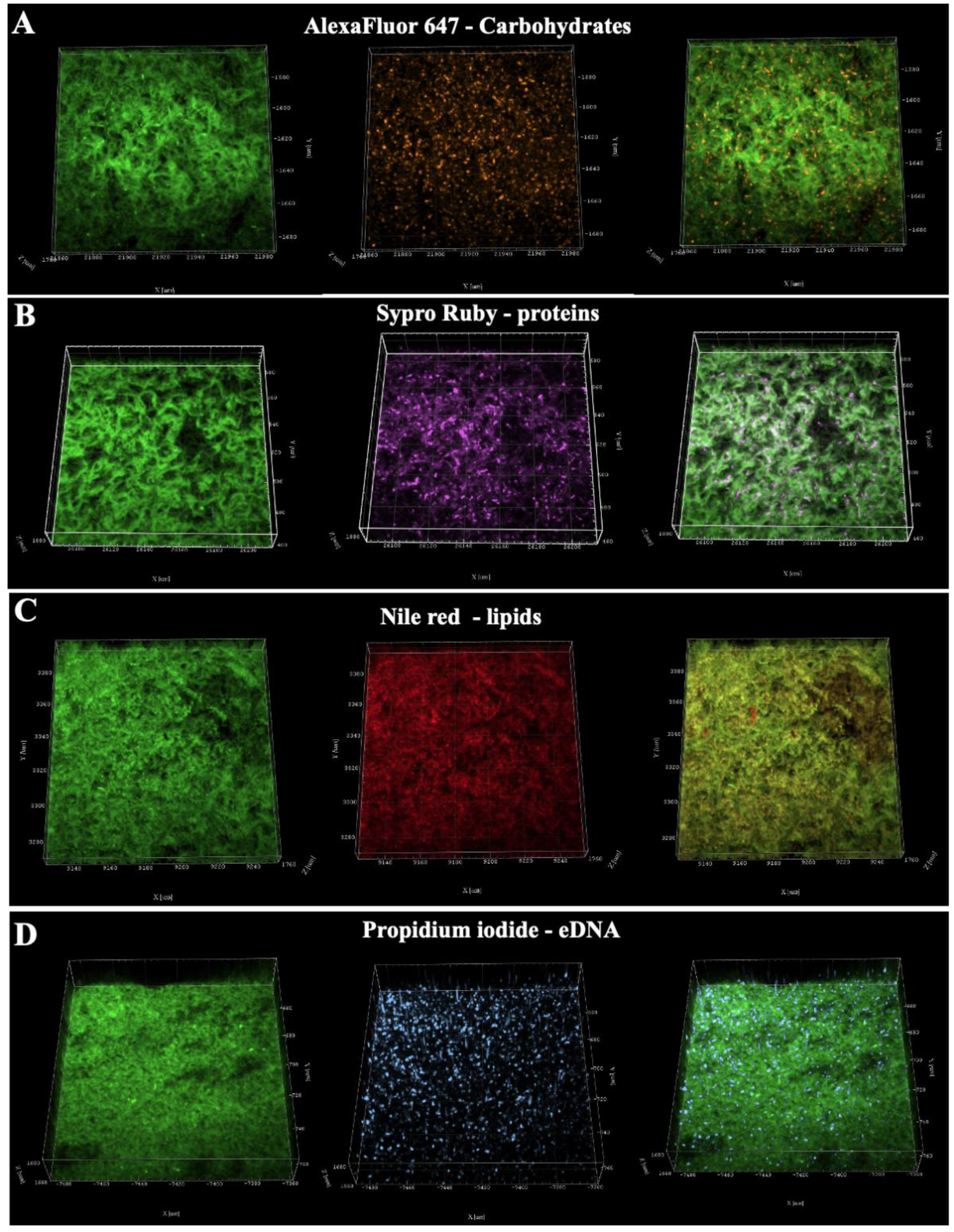
Confocal laser scanning microscopy pictures of 6-day old *M. abscessus* pellicles. 3D projections of confocal z-stacks showing in a row, left to right, eGFP-expressing *M. abscessus*, dyed component of ECM and the overlay; **A**. Carbohydrates stained with Alexa Fluor 647 conjugated to Concanavalin A; **B**. Proteins stained with Sypro Ruby; **C**. Lipids stained with Nile Red; **D**. eDNA stained with Propidium iodide.

Mander’s coefficients measure co-occurrence and were derived as an improvement to PCC since Pearson’s is not sensitive to signal intensity between different parts of an image. M1 and M2 measure the fraction of a given signal that overlaps with another signal. In this case M1 measures how the GFP signal overlaps with a signal emitted from a dye and M2 measures the opposite, how much of the dye signal overlaps with the GFP signal. A value of 1 indicates all of the signal overlaps, while 0 means none of it does. Measuring Mander’s co-efficients is an excellent way to evaluate the co-occurrence of ECM components with cells in biofilms, because it can quantify signal only present in the ECM, meaning the fluorophore signal that does not overlap with the GFP signal. The results of co-localization and biovolume analysis are summarized in **Table 1**.

**Table 1:**
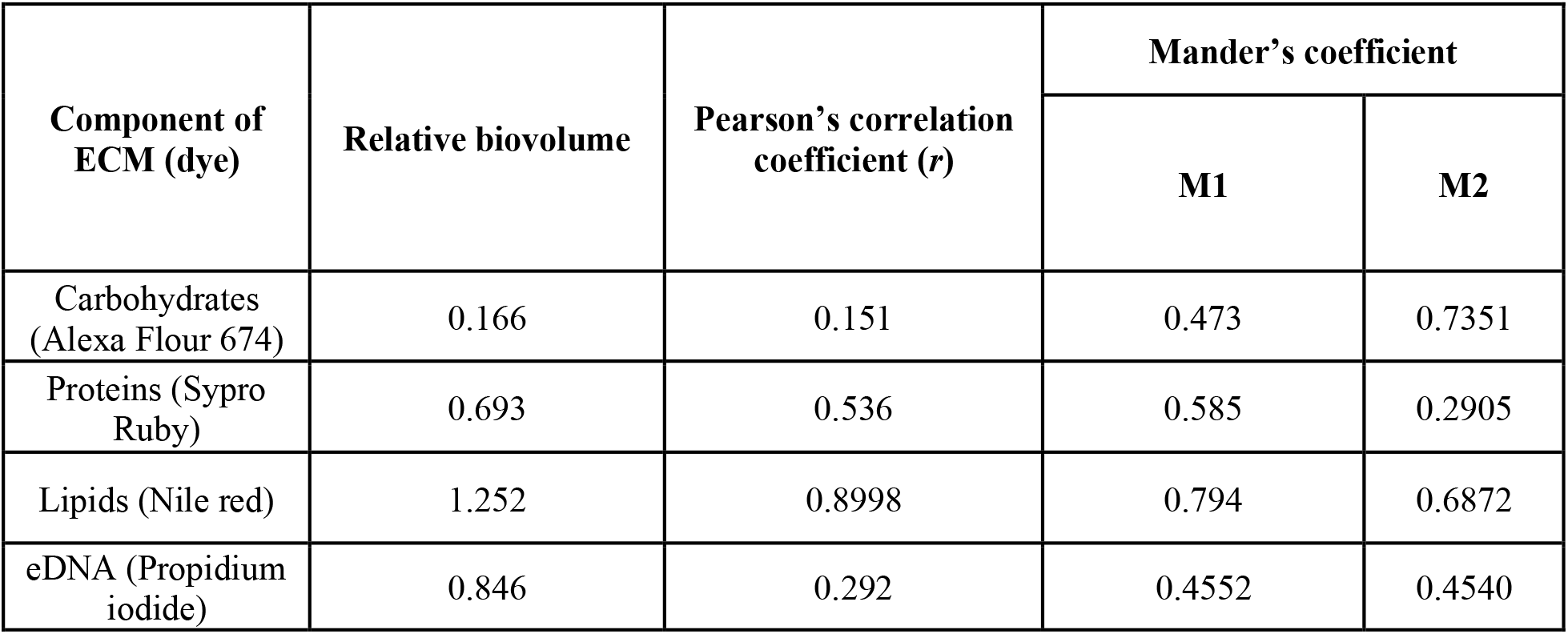
Relative biovolume of ECM components, and colocalization coefficients *r*, M1 and M2 measuring correlation and co-occurrence of eGFP signal with fluorophore signals staining components of the ECM

| Component of ECM (dye) | Relative biovolume | Pearson's correlation coefficient ($r$) | Mander's coefficient | |
| --- | --- | --- | --- | --- |
|  |  |  | M1 | M2 |
| Carbohydrates (Alexa Flour 674) | 0.166 | 0.151 | 0.473 | 0.7351 |
| Proteins (Sypro Ruby) | 0.693 | 0.536 | 0.585 | 0.2905 |
| Lipids (Nile red) | 1.252 | 0.8998 | 0.794 | 0.6872 |
| eDNA (Propidium iodide) | 0.846 | 0.292 | 0.4552 | 0.4540 |

Lipids were the most abundant biomolecule in *M. abscessus* biofilms (relative biovolume=1.252), with a high degree of co-localization with the cells (*r=*0.8998) and a relatively high degree of co-occurrence as indicated by the M1 and M2 (**Table 1**). About 30% of the NR signal did not co-occur with the GFP signal, as indicated by M2=0.6872. This suggested that a fraction of the lipids was found only in the ECM, additionally visible in **File S1, supplementary materials** which shows sectoring of lipids across the ECM and patches of the red signal that do not overlap with green. Proteins, stained with SR, had a medium degree of co-localization (*r=*0.536) with cells, and a low signal overlap of SR with GFP (M2=0.2905), indicating that a high proportion of detected SR signal is coming from proteins present in the ECM, independent of cells. A sector of proteins which did not co-occur with the cells is visible towards the bottom of the binarized z-stack (**File S1, supplementary materials**). The PI signal, staining eDNA, showed poor intensity correlation with GFP (*r*=0.292) and GFP and PI signals overlapped equally with each other at ∼50% (M1=0.4552, M2=0.4540). This suggested that eDNA is dispersed throughout the biofilm, and likely present in areas with low cell count, as seen at the bottom of the biofilm in binarized z-stack images (**File S1, supplementary materials**). Carbohydrates were the smallest component of the *M. abscessus* biofilm (relative biovolume=0.151) and showed a low degree of co-localization (*r=*0.151). Around 70% of the AF signal overlapped with the GFP signal (M2=0.7351) indicating a high degree of co-occurrence, but also showing the remaining ∼30% of AF signal occurred separately from the GFP signal and can be associated with carbohydrates in the ECM, which appear dispersed throughout (**File S1, supplementary materials**). In summary, lipids were the most abundant component of the *M. abscessus* biofilm ECM, while carbohydrates were the least abundant. Some of the fluorescence signals emanating from the dye-stained biomolecules were from distinct sectors with low bacterial presence indicating they are only a part of the ECM.

### 3.3 *M. abscessus* biofilms show differential regulation of lipid metabolism pathways

To further outline potential mechanisms for *M. abscessus* biofilm formation, we used RNA-Seq to sample expression profiles of the biofilms at two key timepoints. The first timepoint, t1, was taken 5 days post inoculation when the biofilm had matured and the second timepoint, t2, was taken 7 days post inoculation, when the biofilm was ready for dispersal. A mid-log planktonic culture was used as a baseline for comparison. Overall, there were 248 significantly differentially expressed genes (DEGs) identified in biofilm t1, (9-down and 239 upregulated) and 1,176 DEGs (513 down- and 663 upregulated) in biofilm t2 (absolute log2 fold-change >2, *P*-value < 0.01). Out of those, 218 were differentially expressed at both timepoints (**Fig. 3A**), had the same directionality (i.e., up- or down-regulated) and were mainly upregulated, indicating commonality in genes likely to be associated with biofilm formation and maintenance. The metabolic pathways associated with the DEGs in biofilm t1 and t2 were assessed using pathway enrichment in comparison to *M. tuberculosis* metabolism from the recently updated genome-scale model iEK1011 (Kavvas et al., 2018). At biofilm maturation, t1, four pathways, related to stress response, fatty acid biosynthesis, glyoxylate biosynthesis and redox metabolism, were upregulated (**Fig. 3C**). Five upregulated genes involved in fatty acid biosynthesis, *MAB_2030, MAB_2028, MAB_2032, MAB_3354* and *MAB_3455c* **(File S2, supplementary materials)** are all involved in mycolic acid chain elongation including a desaturase involved in modification of the meromycolyl chain. At t2, eight pathways showed enrichment: six pathways, related to ABC transporters, quinone metabolism, oxidative phosphorylation, cell division, mammalian cell entry (MCE) and peptidoglycan synthesis, were downregulated, and two pathways, related to oxidoreductase activity and propanoate metabolism, were upregulated (**Fig. 3D**).

**Fig. 3.**
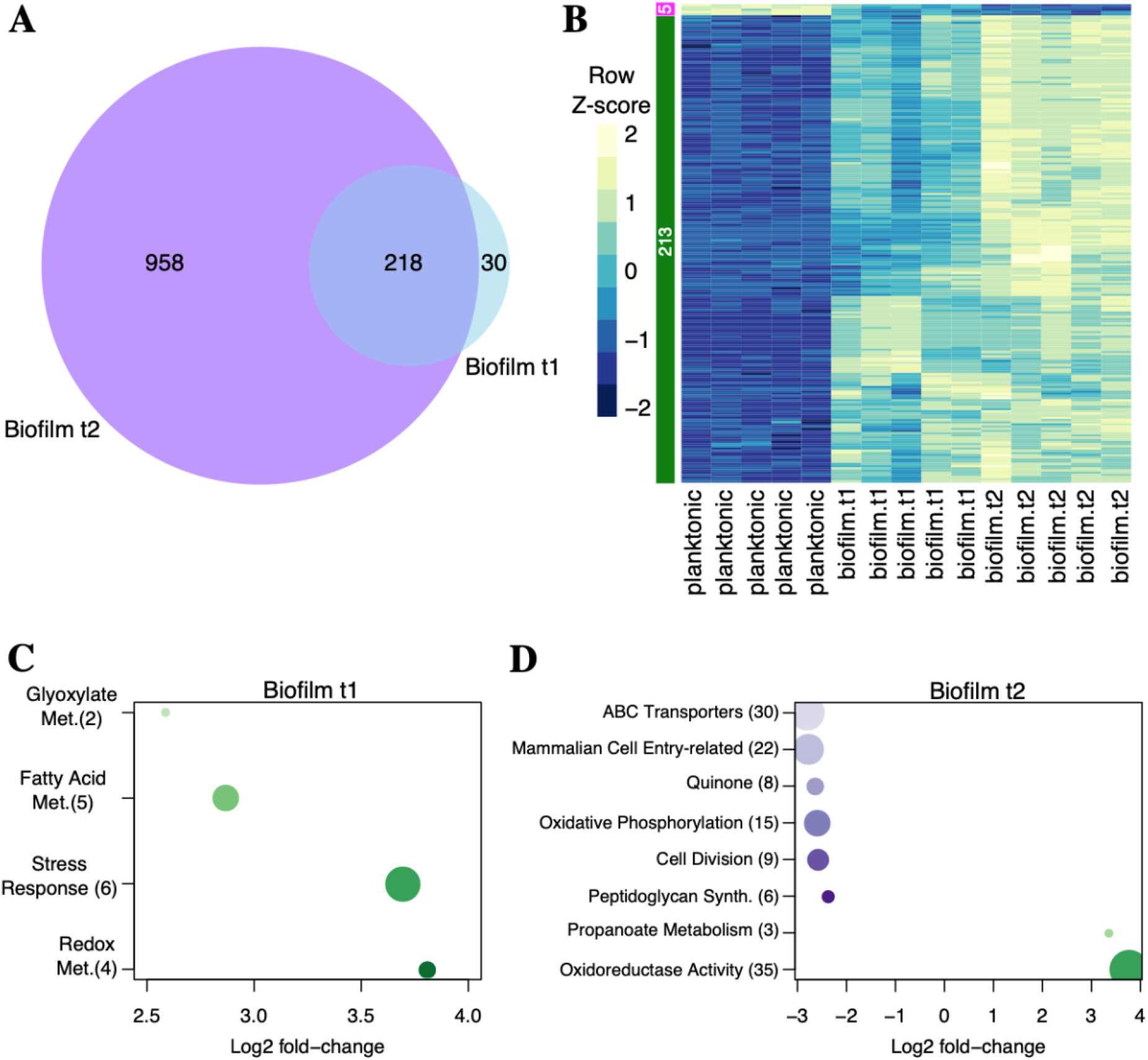
Differentially expressed genes and enriched metabolic pathways in *M. abscessus* biofilms. **A**. Venn diagram illustrating differentially expressed genes (DEGs) during biofilm t1 and t2 with respect to planktonic t1. **B**. Heatmap of transcriptional profiles (z-scores) of the 218 DEGs in biofilm t1 and t2. **C**. and **D**. Lists of enriched metabolic pathways showing log2 fold-change for biofilm t1 and t2 with respect to the planktonic t1 sample. Numbers in parentheses indicate the number of DEGs associated with each functional term. This quantity is also indicated by the size of each circle (the higher the number, the bigger the node). Down- and up- regulated functional terms are indicated with purple and green nodes, respectively. Darker colours indicate higher log2 fold-changes.

### 3.4 Biofilms of *M. abscessus* showed an altered mycolic acid profile

Based on transcription analysis that showed increase in the expression of mycolic acid biosynthesis genes in biofilms, and previous knowledge that lipids are instrumental to biofilm formation in mycobacteria (Ojha et al., 2005b, 2008), we did a comprehensive analysis of lipids in planktonic and biofilm *M. abscessus* (**File S2, supplementary materials**). Lipid fractions extracted from ^14^C supplemented cultures were resolved by thin layer chromatography (TLC) and visualized by autoradiography. The most significant difference in lipid profiles between the two growth conditions was the increase in free mycolic acids in biofilms, a phenotype observed in other mycobacterial biofilms (Ojha et al., 2008; Sophia A Pacheco et al., 2013; Vega-Dominguez et al., 2020) (**Fig. 4A and 4E**). We then extracted mycolic acids as methyl esters (MAMEs) and analysed them by 2D argentated TLC to separate individual mycolic acid subspecies based on degree of desaturation (increased number of double bonds results in retardation in the second AgNO_3_-containing dimension) (**Fig.4 B and C**). We observed a qualitative difference in the patterns of migration of α MAMES. Rapidly migrating α MAMEs, labelled as α_3_* and α_4_* were observed in biofilms (**Fig. 4B and 4C**), and were absent in the planktonic cultures, but replaced by slower migrating species, α_3_ and α_4_. Matrix-assisted laser desorption/ionization-time of flight (MALDI-TOF) analysis of the extracted MAMEs did not show any qualitative differences between biofilm and planktonic cultures, except that those extracted from biofilms were slightly longer compared to those from planktonic cultures (**Fig. 4G and File S4, supplementary materials**). The concurrent replacement of α_3_ and α_4_ MAMEs with the relatively faster migrating α_3_* and α_4_* suggested that α_3_* and α_4_* were likely cyclopropanated derivates of α_3_ and α4 mycolic acids. MALDI-TOF analysis would not reveal the presence of the cyclopropane rings given that that these modified species would simply show a mass reflecting an additional CH_2_ unit. We observed α-mycolic acid species in planktonic cells that range from C-74 to C-79, with the foremost abundant species being C-76. In biofilms we observed a range from C-74 to C-82, with α-79 being the most abundant species, indicating that longer-chain MAMEs were more abundant in biofilms (**Fig. 4G**). From previous studies we know that in α and α’-mycolic acid structure the α-alkyl chain consists of 21 repeating units of CH_2_ with a methyl group at the end, so the variation we see are in the elongation or cyclopropanation of the β-hydroxy chain (Halloum et al., 2016).

**Fig. 4.**
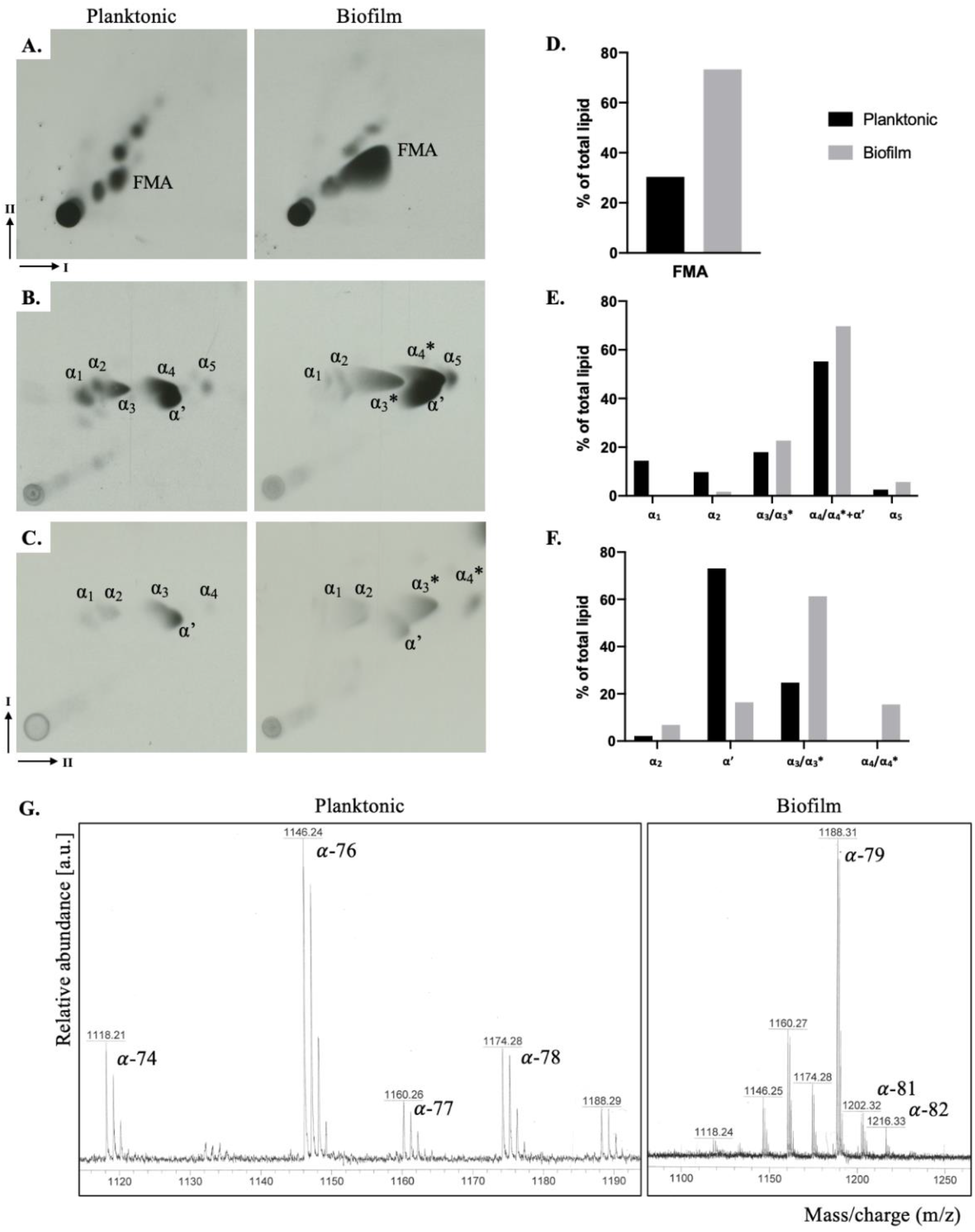
Altered expression of mycolic acids in *M. abscessus* biofilms. Autoradiographs of solvent extractable lipids separated by polarity (**A-C**), with bar graphs showing relative amounts of each labelled spot indicated as a percentage of total amount of lipids detected on the TLC, as determined by densitometry analysis (**D-F**.). **A**. Apolar lipids resolved using solvents in 2D dimensions (I-1^st^, II-2^nd^ dimension), where FMA stands for free mycolic acids; **B**. and **C**. 2D (I-1^st^, II-2^nd^ dimension) argentation TLCs of mycolic acid methyl esters (MAMEs) of wall bound (B.) and apolar lipid fractions (C.). Argentation allows for separation of molecules based on desaturation. α and α’ are mycolic acid subtypes found in *M. abscessus*, *indicates changes in species migration; **G**. MALDI-TOF analysis of wall bound mycolic acids annotated with C chain lengths that each molecular weight represents, more detail can be found in File S4, supplementary materials.

## 4 Discussion

In this study we characterized the *in vitro* formation of a pellicular biofilm in *M. abscessus*, describing the appearance and contents of the ECM and determining its unique transcriptional profile with a particular focus on the impact on changes in mycolic acid expression. We visualized the ubiquitous pattern of cellular organization, where cells enveloped in an ECM were aggregated in an organized manner forming channels and pores that are also visible through the ECM (**Fig. 1**). Using CLSM, we showed that the ECM consists of lipids, polysaccharides, proteins and eDNA (**Fig. 2**), all previously reported to play integral roles in biofilm formation and maturation in various bacterial species (Rose et al., 2015). We found that eDNA is abundant in *M. abscessus* biofilms (relative biovolume=0.846) and dispersed throughout the biofilm, particularly in areas with low cell count which points to eDNA having a role in aggregation of the biofilm mirroring its function in surface-attached biofilms, previously outlined in *M. avium* studies (Rose et al., 2015). In contrast to the significant role carbohydrates play in *M. chelonae* and *M. tuberculosis* (Chakraborty and Kumar, 2019; Trivedi et al., 2016; Vega-Dominguez et al., 2020), in *M. abscessus* biofilms carbohydrates appear dispersed as evidenced by the trace presence in ECM (**Table 1**). A study by Lemassu *et al*., 1996 similarly showed that surface exposed material of RGM is protein-rich rather than carbohydrate-rich, in direct contrast to SGM where the opposite is true – a correlation to the differences in the amount of carbohydrates reported in *M. chelonae* and *M. tuberculosis* and our reports in *M. abscessus*.

In *M. abscessus* biofilms lipids were found to co-localize with cells to a high degree (*r*=0.8998), however Mander’s coefficients showed that while a significant portion of the NR signal co-occurred with GFP signal (M2=0.687), and vice versa (M1=0.794), a significant portion of the signal did not overlap indicating presence of lipids in the ECM separately from the cells (**Table 1, Fig.2C**). As a result of the unique make up of mycobacterial membranes, function of lipids has been thoroughly researched and several species are known constituents of the ECM, or have been shown to play an important role in biofilm formation and maintenance (Ripoll et al., 2007; Schorey and Sweet, 2008; Trivedi et al., 2016; Yamazaki et al., 2006). Deletion of genes responsible for glycopeptidolipids biosynthesis impaired biofilm formation in *M. avium* (Yamazaki et al., 2006), however analysis showed no notable difference in GPL expression between planktonic and biofilm growth in *M. abscessus* (data not shown) (Ripoll et al., 2007; Schorey and Sweet, 2008). An increase in phosphatidyl myo-inositol mannosides (PIMs), however, was noted (**File S3, supplementary materials**), which could point to a role in maintenance of biofilm structure, since altered acetylation of PIMs was shown to lead to defective biofilm formation (Li et al., 2020).

A notable outcome of this study is the visible increase in free mycolic acid in biofilms (**Fig. 4**), often described a hallmark of biofilm formation in NTMs (Ojha et al., 2005a, 2008; Sambandan et al., 2013; Vega-Dominguez et al., 2020). α and α’-mycolic acid are the two identified mycolic acid subtypes in *M. abscessus* planktonic cells (Halloum et al., 2016). Alpha-mycolic acid consists of a long carbon chain (C_74_-C_79_) containing two *cis/trans* double bonds or *cis*-cyclopropyl groups in the meromycolate chain, while α’-mycolic acid has a shorter carbon chain (C_62_-C_64_) and contains only a single *cis* double bond (Marrakchi et al., 2014). We report an accumulation of free mycolic acid in biofilms (**Fig 4.A**), and a change in α-mycolic acid indicative of an increase in chain length and a decrease in desaturation, or more likely an increase in cyclopropanation. Importance of longer chain mycolic acids in biofilms was showed through studies in *M. smegmatis* and *M. bovis* BCG where the deletion of enzymes related to mycolic acid biosynthesis, specifically that encode KasB and GroEL1, a KasA chaperone, caused a decrease in mycolate chain length and impacted biofilm formation (Gao et al., 2003; Ojha et al., 2005b; Wang et al., 2011). In *M. abscessus* biofilms we report an upregulation of the mycolic acid pathway (**Fig. 3**), with a specific focus on elongation and desaturation of the meromycolic chain. There is a strong upregulation of *MAB_2030* and *MAB_2028*, encoding for KasA/B complex which is responsible for elongation and *MAB_3354*, encoding for DesA1, a desaturase protein responsible for addition of double bonds to the β-hydroxy branch (A. Singh et al., 2016). The evidence from our phenotypic study, coupled with the transcriptional analysis, shows that in *M. abscessus* biofilms there is an accumulation of long chained mycolic acids, with an initial increase in desaturated molecules that likely become cyclopropanated.

Upregulation of the enzymes of the glyoxylate shunt, as seen in *M. abscessus* biofilm t1 (**Fig. 3C**), was previously reported in nutrient starvation, hypoxia, and in *M. tuberculosis* persister cells (Giffin et al., 2016; Miranda-CasoLuengo et al., 2016). Isocitrate lyase, *MAB_4095c*, which is upregulated as part of this pathway, is necessary for establishment of chronic infections with *M. tuberculosis* (McKinney et al., 2000), and inhibition of this enzyme forces bacteria to re-enter a replicative stage, implying its importance in dormancy and biofilms (Van Schaik et al., 2009). Together the upregulation of glyoxylate and fatty acid metabolism was shown to allow bacteria to survive in low nutrient environments in biofilms and to modify the cell surface and decrease susceptibility to antibiotics (Rojony et al., 2020, 2019), an event we are likely registering in *M. abscessus* biofilms as well. Upregulation of the redox metabolism and stress in biofilm t1 (**Fig. 3C**), and oxidoreductases in biofilm t2, indicate an adaptation of the cells to the new biofilm environment, or an indication that certain parts of the biofilm are not receiving enough oxygen. In *M. smegmatis* the ratio of NADH/NAD^+^ is three-fold higher in biofilm than in planktonic cells representative of a reductive environment, and fluctuations in the redox state of the bacilli and (p)ppGpp and c-di-GMP, markers of stress response, play critical roles in biofilm formation (Geier et al., 2008; Gupta et al., 2015; Trivedi et al., 2016; Wolff et al., 2015). Upregulated genes in these pathways were mainly encode for subunits of the cytochrome D ubiquinol oxidases, CydA, CydB and CydD, all shown to be necessary in *E. coli* to form a mature biofilm and an ECM (Beebout et al., 2019). Additionally we found genes responsible for mammalian cell entry (MCE) to be downregulated in the biofilm late stage (t2) (**Fig. 3**D) similarly to reports in *M. chelonae* biofilms (Vega-Dominguez et al., 2020) but contrary to those *M. tuberculosis* where *mce* operons are upregulated in stationary phase (P. Singh et al., 2016). Downregulation of genes encoding for MCE and for ABC transporters could be linked to the increase in mycolic acids in *M. abscessus* biofilms (**Fig. 4A**), given that *mce1* encodes 13 genes responsible for lipid transport and that the accumulation of mycolic acids was previously seen in Δ*mce1 M. tuberculosis* (Cantrell et al., 2013).

In conclusion, the transition from planktonic to biofilm growth in *M. abscessus* induces a significant metabolic change reflected in the phenotype. *M. abscessus* biofilms ECM is composed of carbohydrates, lipids, proteins and eDNA, with lipids making up the biggest component and MAs specifically undergoing molecular changes traceable in the transcription profile. Upregulation of genes that encode for elongation and desaturation of α-mycolates is reflected in the shift to longer carbon chains in biofilm MAs. The upregulation in glyoxylate metabolism and isocitrate lyase, and the downregulation in cell division and peptidoglycan biosynthesis in biofilms are indicative of an environment of energy preservation, while the increase in redox metabolism and oxidoreductases suggest an anaerobic environment in parts of the biofilm. In fact, the CF lung environment, in which *M. abscessus* forms biofilm-like aggregates, is known to have hypoxic conditions and it was shown that *in vitro* bacterial aggregation leads to a decrease in O_2_ consumption leading to slow growth, low aerobic respiration and an increased resistance to aminoglycosides (Kolpen et al., 2020). Overall, molecular changes that occur in the *M. abscessus* biofilm promote survival, protection and modulation of the cell envelope. This initial characterization of the *M. abscessus* pellicle gives a firm steppingstone for further studies into clinical treatments for dispersion of *M. abscessus* biofilms.

## Acknowledgments

AD was supported by a PhD studentship from the Darwin Trust of Edinburgh. We also thank Laurent Kremer for providing the plasmid pMV306eGFP. AB acknowledges funding from the Medical Research Council (MR/S000542/1) and the BBSRC-NSF (BB/N01314X/1). Funding was also provided by the National Science Foundation [1518261] and the National Institutes of Health [R01AI12825 and R01AI141953].

## Data availability

The RNA-seq data (raw fastq files and read counts) have been deposited in the GEO repository under accession number GSE165352. The authors declare that all relevant data supporting the findings of the study are available in this article and the Supplementary materials, or from the corresponding author upon reasonable request.

## References

Aung, T.T., Chor, W.H.J., Yam, J.K.H., Givskov, M., Yang, L., Beuerman, R.W., 2017. Discovery of novel antimycobacterial drug therapy in biofilm of pathogenic nontuberculous mycobacterial keratitis. Ocul. Surf. https://doi.org/10.1016/j.jtos.2017.06.002

Aung, T.T., Yam, J.K.H., Lin, S., Salleh, S.M., Givskov, M., Liu, S., Lwin, N.C., Yang, L., Beuerman, R.W., 2016. Biofilms of pathogenic nontuberculous mycobacteria targeted by new therapeutic approaches. Antimicrob. Agents Chemother. https://doi.org/10.1128/AAC.01509-15

Beebout, C.J., Eberly, A.R., Werby, S.H., Reasoner, S.A., Brannon, J.R., De, S., Fitzgerald, M.J., Huggins, M.M., Clayton, D.B., Cegelski, L., Hadjifrangiskou, M., 2019. Respiratory heterogeneity shapes biofilm formation and host colonization in uropathogenic Escherichia coli. MBio. https://doi.org/10.1128/mBio.02400-18

Besra, G.S., 1998. Preparation of Cell-Wall Fractions from Mycobacteria, in: Mycobacteria Protocols. Humana Press, New Jersey, pp. 91–104. https://doi.org/10.1385/0-89603-471-2:91

Bosio, S., Leekha, S., Gamb, S.I., Wright, A.J., Terrell, C.L., Miller, D. V., 2012. Mycobacterium fortuitum prosthetic valve endocarditis: a case for the pathogenetic role of biofilms. Cardiovasc. Pathol. 21, 361–364. https://doi.org/10.1016/j.carpath.2011.11.001

Cantrell, S.A., Leavell, M.D., Marjanovic, O., Iavarone, A.T., Leary, J.A., Riley, L.W., 2013. Free mycolic acid accumulation in the cell wall of the mce1 operon mutant strain of Mycobacterium tuberculosis. J. Microbiol. https://doi.org/10.1007/s12275-013-3092-y

Cavaliere, R., Ball, J.L., Turnbull, L., Whitchurch, C.B., 2014. The biofilm matrix destabilizers, EDTA and DNaseI, enhance the susceptibility of nontypeable Hemophilus influenzae biofilms to treatment with ampicillin and ciprofloxacin. Microbiologyopen 3, 557–567. https://doi.org/10.1002/mbo3.187

Chakraborty, P., Kumar, A., 2019. The extracellular matrix of mycobacterial biofilms: Could we shorten the treatment of mycobacterial infections? Microb. Cell. https://doi.org/10.15698/mic2019.02.667

Chen, J.M., German, G.J., Alexander, D.C., Ren, H., Tan, T., Liu, J., 2006. Roles of Lsr2 in colony morphology and biofilm formation of Mycobacterium smegmatis. J. Bacteriol. https://doi.org/10.1128/JB.188.2.633-641.2006

Cystic Fibrosis Foundation, 2010. Cystic Fibrosis Foundation Patient Registry, 2010 Annual Data Report. Cyst. Fibros. Found. Publ.

De Chaumont, F., Dallongeville, S., Chenouard, N., Hervé, N., Pop, S., Provoost, T., et al., 2012. Icy: an open bioimage informatics platform for extended reproducible research. Nat. Methods 9, 690–6.

Faria, S., Joao, I., Jordao, L., 2015. General Overview on Nontuberculous Mycobacteria, Biofilms, and Human Infection. J. Pathog. 2015, 1–10. https://doi.org/10.1155/2015/809014

Fennelly, K.P., Ojano-Dirain, C., Yang, Q., Liu, L., Lu, L., Progulske-Fox, A., Wang, G.P., Antonelli, P., Schultz, G., 2016. Biofilm Formation by Mycobacterium abscessus in a Lung Cavity. Am. J. Respir. Crit. Care Med. 193, 692–693. https://doi.org/10.1164/rccm.201508-1586IM

Flemming, H.-C., Wingender, J., 2010. The biofilm matrix. Nat. Publ. Gr. 8. https://doi.org/10.1038/nrmicro2415

Flemming, H.C., Wingender, J., Szewzyk, U., Steinberg, P., Rice, S.A., Kjelleberg, S., 2016. Biofilms: An emergent form of bacterial life. Nat. Rev. Microbiol. https://doi.org/10.1038/nrmicro.2016.94

Galassi, L., Donato, R., Tortoli, E., Burrini, D., Santianni, D., Dei, R., 2003. Nontuberculous mycobacteria in hospital water systems: Application of HPLC for identification of environmental mycobacteria. J. Water Health 1, 133–139.

Gao, L.Y., Laval, F., Lawson, E.H., Groger, R.K., Woodruff, A., Morisaki, J.H., Cox, J.S., Daffe, M., Brown, E.J., 2003. Requirement for kasB in Mycobacterium mycolic acid biosynthesis, cell wall impermeability and intracellular survival: Implications for therapy. Mol. Microbiol. https://doi.org/10.1046/j.1365-2958.2003.03667.x

Geier, H., Mostowy, S., Cangelosi, G.A., Behr, M.A., Ford, T.E., 2008. Autoinducer-2 triggers the oxidative stress response in Mycobacterium avium, leading to biofilm formation. Appl. Environ. Microbiol. https://doi.org/10.1128/AEM.02066-07

Ghosh, R., Das, S., Kela, H., De, A., Haldar, J., Maiti, P.K., 2017. Biofilm colonization of Mycobacterium abscessus: New threat in hospital-acquired surgical site infection. Indian J. Tuberc. 64, 178–182. https://doi.org/10.1016/J.IJTB.2016.11.013

Giffin, M.M., Shi, L., Gennaro, M.L., Sohaskey, C.D., 2016. Role of Alanine Dehydrogenase of Mycobacterium tuberculosis during Recovery from Hypoxic Nonreplicating Persistence. PLoS One. https://doi.org/10.1371/journal.pone.0155522

Greendyke, R., Byrd, T.F., 2008. Differential antibiotic susceptibility of Mycobacterium abscessus variants in biofilms and macrophages compared to that of planktonic bacteria. Antimicrob. Agents Chemother. 52, 2019–2026. https://doi.org/10.1128/AAC.00986-07

Gupta, K.R., Kasetty, S., Chatterji, D., 2015. Novel functions of (p)ppGpp and cyclic di-GMP in mycobacterial physiology revealed by phenotype microarray analysis of wild-type and isogenic strains of Mycobacterium smegmatis. Appl. Environ. Microbiol. https://doi.org/10.1128/AEM.03999-14

Halloum, I., Carrère-Kremer, S., Blaise, M., Viljoen, A., Bernut, A., Le Moigne, V., Vilchèze, C., Guérardel, Y., Lutfalla, G., Herrmann, J.-L., Jacobs, W.R., Kremer, L., Kremer, L., 2016. Deletion of a dehydratase important for intracellular growth and cording renders rough Mycobacterium abscessus avirulent. Proc. Natl. Acad. Sci. U. S. A. 113, E4228–37. https://doi.org/10.1073/pnas.1605477113

Howard, S.T., Rhoades, E., Recht, J., Pang, X., Alsup, A., Kolter, R., Lyons, C.R., Byrd, T.F., 2006. Spontaneous reversion of Mycobacterium abscessus from a smooth to a rough morphotype is associated with reduced expression of glycopeptidolipid and reacquisition of an invasive phenotype. Microbiology 152, 1581–1590. https://doi.org/10.1099/mic.0.28625-0

Ibáñez de Aldecoa, A.L., Zafra, O., González-Pastor, J.E., 2017. Mechanisms and regulation of extracellular DNA release and its biological roles in microbial communities. Front. Microbiol. https://doi.org/10.3389/fmicb.2017.01390

Karygianni, L., Ren, Z., Koo, H., Thurnheer, T., 2020. Biofilm Matrixome: Extracellular Components in Structured Microbial Communities. Trends Microbiol. https://doi.org/10.1016/j.tim.2020.03.016

Kavvas, E.S., Seif, Y., Yurkovich, J.T., Norsigian, C., Poudel, S., Greenwald, W.W., Ghatak, S., Palsson, B.O., Monk, J.M., 2018. Updated and standardized genome-scale reconstruction of Mycobacterium tuberculosis H37Rv, iEK1011, simulates flux states indicative of physiological conditions. BMC Syst. Biol. https://doi.org/10.1186/s12918-018-0557-y

Kim, S.-Y., Shin, S.J., Jeong, B.-H., Koh, W.-J., 2016. Successful antibiotic treatment of pulmonary disease caused by Mycobacterium abscessus subsp. abscessus with C-to-T mutation at position 19 in erm(41) gene: case report. BMC Infect. Dis. 16, 207. https://doi.org/10.1186/s12879-016-1554-7

Kolpen, M., Jensen, P.Ø., Qvist, T., Kragh, K.N., Ravnholt, C., Fritz, B.G., Johansen, U.R., Bjarnsholt, T., Hoiby, N., 2020. Biofilms of mycobacterium abscessus complex can be sensitized to antibiotics by disaggregation and oxygenation. Antimicrob. Agents Chemother. https://doi.org/10.1128/AAC.01212-19

Kwak, N., Dalcolmo, M.P., Daley, C.L., Eather, G., Gayoso, R., Hasegawa, N., Jhun, B.W., Koh, W.J., Namkoong, H., Park, J., Thomson, R., Van Ingen, J., Zweijpfenning, S.M.H., Yim, J.J., 2019. Mycobacterium abscessus pulmonary disease: Individual patient data meta-analysis. Eur. Respir. J. https://doi.org/10.1183/13993003.01991-2018

Lagache, T., Sauvonnet, N., Danglot, L., Olivo-Marin, J.C., 2015. Statistical analysis of molecule colocalization in bioimaging. Cytom. Part A. https://doi.org/10.1002/cyto.a.22629

Lemassu, A., Ortalo-Magne, A., Bardou, F., Silve, G., Laneelle, M.-A., Daffe, M., Ortalo-Magné, A., Bardou, F., Silve, G., Lanéelle, M.A., Daffé, M., 1996a. Extracellular and surface-exposed polysaccharides of non-tuberculous mycobacteria. Microbiology 142, 1513–1520. https://doi.org/10.1099/13500872-142-6-1513

Lemassu, A., Ortalo-Magné, A., Bardou, F., Silve, G., Lanéelle, M.A., Daffé, M., 1996b. Extracellular and surface-exposed polysaccharides of non-tuberculous mycobacteria. Microbiology. https://doi.org/10.1099/13500872-142-6-1513

Li, M., Gašparovič, H., Weng, X., Chen, S., Korduláková, J., Jessen-Trefzer, C., 2020. The Two-Component Locus MSMEG_0244/0246 Together With MSMEG_0243 Affects Biofilm Assembly in M. smegmatis Correlating With Changes in Phosphatidylinositol Mannosides Acylation. Front. Microbiol. 11. https://doi.org/10.3389/fmicb.2020.570606

Marrakchi, H., Lanéelle, M.A., Daffé, M., 2014. Mycolic acids: Structures, biosynthesis, and beyond. Chem. Biol. https://doi.org/10.1016/j.chembiol.2013.11.011

McKinney, J.D., Höner Zu Bentrup, K., Muñoz-Elias, E.J., Miczak, A., Chen, B., Chan, W.T., Swenson, D., Sacchettini, J.C., Jacobs, W.R., Russell, D.G., 2000. Persistence of Mycobacterium tuberculosis in macrophages and mice requires the glyoxylate shunt enzyme isocitrate lyase. Nature. https://doi.org/10.1038/35021074

Medjahed, H., Gaillard, J.L., Reyrat, J.M., 2010. Mycobacterium abscessus: a new player in the mycobacterial field. Trends Microbiol. https://doi.org/10.1016/j.tim.2009.12.007

Miranda-CasoLuengo, A.A., Staunton, P.M., Dinan, A.M., Lohan, A.J., Loftus, B.J., 2016. Functional characterization of the Mycobacterium abscessus genome coupled with condition specific transcriptomics reveals conserved molecular strategies for host adaptation and persistence. BMC Genomics 17, 553. https://doi.org/10.1186/s12864-016-2868-y

Ojha, A., Anand, M., Bhatt, A., Kremer, L., Jacobs, W.R., Hatfull, G.F., 2005a. GroEL1: A dedicated chaperone involved in mycolic acid biosynthesis during biofilm formation in mycobacteria. Cell. https://doi.org/10.1016/j.cell.2005.09.012

Ojha, A., Anand, M., Bhatt, A., Kremer, L., Jacobs, W.R., Hatfull, G.F., 2005b. GroEL1: A dedicated chaperone involved in mycolic acid biosynthesis during biofilm formation in mycobacteria. Cell 123, 861–873. https://doi.org/10.1016/j.cell.2005.09.012

Ojha, A.K., Baughn, A.D., Sambandan, D., Hsu, T., Trivelli, X., Guerardel, Y., Alahari, A., Kremer, L., Jacobs, W.R., Hatfull, G.F., 2008. Growth of Mycobacterium tuberculosis biofilms containing free mycolic acids and harbouring drug-tolerant bacteria. Mol. Microbiol. 69, 164–174. https://doi.org/10.1111/j.1365-2958.2008.06274.x

Pacheco, Sophia A, Hsu, F.-F., Powers, K.M., Purdy, G.E., 2013. MmpL11 protein transports mycolic acid-containing lipids to the mycobacterial cell wall and contributes to biofilm formation in Mycobacterium smegmatis. J. Biol. Chem. 288, 24213–22. https://doi.org/10.1074/jbc.M113.473371

Pacheco, Sophia A., Hsu, F.-F.F., Powers, K.M., Purdy, G.E., 2013. MmpL11 protein transports mycolic acid-containing lipids to the mycobacterial cell wall and contributes to biofilm formation in Mycobacterium smegmatis. J. Biol. Chem. 288, 24213–24222. https://doi.org/10.1074/jbc.M113.473371

Padilla, P., Ly, P., Dillard, R., Boukovalas, S., Zapata-Sirvent, R., Phillips, L.G., 2018. Medical tourism and postoperative infections: A systematic literature review of causative organisms and empiric treatment. Plast. Reconstr. Surg. https://doi.org/10.1097/PRS.0000000000005014

Pakkulnan, R., Anutrakunchai, C., Kanthawong, S., Taweechaisupapong, S., Chareonsudjai, P., Chareonsudjai, S., 2019. Extracellular DNA facilitates bacterial adhesion during Burkholderia pseudomallei biofilm formation. PLoS One. https://doi.org/10.1371/journal.pone.0213288

Pike, J.A., Styles, I.B., Rappoport, J.Z., Heath, J.K., 2017. Quantifying receptor trafficking and colocalization with confocal microscopy. Methods. https://doi.org/10.1016/j.ymeth.2017.01.005

Qvist, T., Eickhardt-Sørensen, S.R., Katzenstein, T.L., Pressler, T., Iversen, M., Andersen, C.B., Bjarnsholt, T., Høiby, N., 2013. WS1.5 First evidence of Mycobacterium abscessus biofilm in the lungs of chronically infected CF patients. J. Cyst. Fibros. 12, S2. https://doi.org/10.1016/S1569-1993(13)60006-5

Qvist, T., Eickhardt, S., Kragh, K.N., Andersen, C.B., Iversen, M., H??iby, N., Bjarnsholt, T., Høiby, N., Bjarnsholt, T., 2015. Chronic pulmonary disease with Mycobacterium abscessus complex is a biofilm infection. Eur. Respir. J. 46, 1823–1826. https://doi.org/10.1183/13993003.01102-2015

Ratnatunga, C.N., Lutzky, V.P., Kupz, A., Doolan, D.L., Reid, D.W., Field, M., Bell, S.C., Thomson, R.M., Miles, J.J., 2020. The Rise of Non-Tuberculosis Mycobacterial Lung Disease. Front. Immunol. https://doi.org/10.3389/fimmu.2020.00303

Recht, J., Kolter, R., 2001. Glycopeptidolipid acetylation affects sliding motility and biofilm formation in Mycobacterium smegmatis. J. Bacteriol. 183, 5718–24. https://doi.org/10.1128/JB.183.19.5718-5724.2001

Rhoades, E.R., Archambault, A.S., Greendyke, R., Hsu, F.-F., Streeter, C., Byrd, T.F., 2009. Mycobacterium abscessus Glycopeptidolipids Mask Underlying Cell Wall Phosphatidyl-myo -Inositol Mannosides Blocking Induction of Human Macrophage TNF-α by Preventing Interaction with TLR2. J. Immunol. https://doi.org/10.4049/jimmunol.0802181

Ripoll, F., Deshayes, C., Pasek, S., Laval, F., Beretti, J.-L., Biet, F., Risler, J.-L., Daffé, M., Etienne, G., Gaillard, J.-L., Reyrat, J.-M., 2007. Genomics of glycopeptidolipid biosynthesis in Mycobacterium abscessus and M. chelonae. BMC Genomics 8, 114. https://doi.org/10.1186/1471-2164-8-114

Rojony, R., Danelishvili, L., Campeau, A., Wozniak, J.M., Gonzalez, D.J., Bermudez, L.E., 2020. Exposure of mycobacterium abscessus to environmental stress and clinically used antibiotics reveals common proteome response among pathogenic mycobacteria. Microorganisms. https://doi.org/10.3390/microorganisms8050698

Rojony, R., Martin, M., Campeau, A., Wozniak, J.M., Gonzalez, D.J., Jaiswal, P., Danelishvili, L., Bermudez, L.E., 2019. Quantitative analysis of Mycobacterium avium subsp. hominissuis proteome in response to antibiotics and during exposure to different environmental conditions. Clin. Proteomics. https://doi.org/10.1186/s12014-019-9260-2

Rose, S.J., Babrak, L.M., Bermudez, L.E. LE, Bermudez, L.E. LE, Kalscheuer, R., Veeraraghavan, U., 2015. Mycobacterium avium Possesses Extracellular DNA that Contributes to Biofilm Formation, Structural Integrity, and Tolerance to Antibiotics. PLoS One 10, e0128772. https://doi.org/10.1371/journal.pone.0128772

Rose, S.J., Bermudez, L.E., 2016. Identification of Bicarbonate as a Trigger and Genes Involved with Extracellular DNA Export in Mycobacterial Biofilms. MBio 7, e01597–16. https://doi.org/10.1128/mBio.01597-16

Roux, A.-L., Viljoen, A., Bah, A., Simeone, R., Bernut, A., Laencina, L., Deramaudt, T., Rottman, M., Gaillard, J.-L., Majlessi, L., Brosch, R., Girard-Misguich, F., Vergne, I., de Chastellier, C., Kremer, L., Herrmann, J.-L., 2016. The distinct fate of smooth and rough Mycobacterium abscessus variants inside macrophages. Open Biol. 6, 160185. https://doi.org/10.1098/rsob.160185

Rüegg, E., Cheretakis, A., Modarressi, A., Harbarth, S., Pittet-Cuénod, B., 2015. Multisite Infection with Mycobacterium abscessus after Replacement of Breast Implants and Gluteal Lipofilling. Case Rep. Infect. Dis. 2015, 1–6. https://doi.org/10.1155/2015/361340

Sambandan, D., Dao, D.N., Weinrick, B.C., Vilchèze, C., Gurcha, S.S., Ojha, A., Kremer, L., Besra, G.S., Hatfull, G.F., Jacobs, W.R., Jr, 2013. Keto-mycolic acid-dependent pellicle formation confers tolerance to drug-sensitive Mycobacterium tuberculosis. MBio 4, e00222–13. https://doi.org/10.1128/mBio.00222-13

Schlafer, S., Meyer, R.L., 2017. Confocal microscopy imaging of the biofilm matrix. J. Microbiol. Methods. https://doi.org/10.1016/j.mimet.2016.03.002

Schorey, J.S., Sweet, L., 2008. The mycobacterial glycopeptidolipids: Structure, function, and their role in pathogenesis. Glycobiology. https://doi.org/10.1093/glycob/cwn076

Singh, A., Varela, C., Bhatt, K., Veerapen, N., Lee, O.Y.C.C., Wu, H.H.T.T., Besra, G.S., Minnikin, D.E., Fujiwara, N., Teramoto, K., Bhatt, A., 2016. Identification of a Desaturase Involved in Mycolic Acid Biosynthesis in Mycobacterium smegmatis. PLoS One 11, e0164253. https://doi.org/10.1371/journal.pone.0164253

Singh, P., Katoch, V.M., Mohanty, K.K., Chauhan, D.S., 2016. Analysis of expression profile of mce operon genes (mce1, mce2, mce3 operon) in different Mycobacterium tuberculosis isolates at different growth phases. Indian J. Med. Res. https://doi.org/10.4103/0971-5916.184305

Solokhina, A., Brückner, D., Bonkat, G., Braissant, O., 2017. Metabolic activity of mature biofilms of Mycobacterium tuberculosis and other non-tuberculous mycobacteria. Sci. Rep. 7, 9225. https://doi.org/10.1038/s41598-017-10019-4

Sousa, S., Bandeira, M., Carvalho, P.A., Duarte, A., Jordao, L., 2015. Nontuberculous mycobacteria pathogenesis and biofilm assembly. Int. J. Mycobacteriology 4, 36–43.

Tetz, G. V., Artemenko, N.K., Tetz, V. V., 2009. Effect of DNase and antibiotics on biofilm characteristics. Antimicrob. Agents Chemother. https://doi.org/10.1128/AAC.00471-08

Toyofuku, M., Inaba, T., Kiyokawa, T., Obana, N., Yawata, Y., Nomura, N., 2016. Environmental factors that shape biofilm formation. Biosci. Biotechnol. Biochem. 80, 7– 12. https://doi.org/10.1080/09168451.2015.1058701

Trivedi, A., Mavi, P.S., Bhatt, D., Kumar, A., 2016. Thiol reductive stress induces cellulose-anchored biofilm formation in Mycobacterium tuberculosis. Nat. Commun. 7, 11392. https://doi.org/10.1038/ncomms11392

Van Schaik, E.J., Tom, M., Woods, D.E., 2009. Burkholderia pseudomallei isocitrate lyase is a persistence factor in pulmonary melioidosis: Implications for the development of isocitrate lyase inhibitors as novel antimicrobials. Infect. Immun. https://doi.org/10.1128/IAI.00609-09

Vasudevan, R., 2014. Biofilms: Microbial Cities of Scientific Significance. J. Microbiol. Exp. 1, 1–0.

Vega-Dominguez, P., Peterson, E., Pan, M., Di Maio, A., Singh, S., Umapathy, S., Saini, D.K., Baliga, N., Bhatt, A., 2020. Biofilms of the non-tuberculous Mycobacterium chelonae form an extracellular matrix and display distinct expression patterns. Cell Surf. https://doi.org/10.1016/j.tcsw.2020.100043

Wang, X.M., Lu, C., Soetaert, K., Heeren, C.S., Peirs, P., Lanéelle, M.A., Lefèvre, P., Bifani, P., Content, J., Daffé, M., Huygen, K., de Bruyn, J., Wattiez, R., 2011. Biochemical and immunological characterization of a cpn60.1 knockout mutant of Mycobacterium bovis BCG. Microbiology. https://doi.org/10.1099/mic.0.045120-0

Whitchurch, C.B., Tolker-Nielsen, T., Ragas, P.C., Mattick, J.S., 2002. Extracellular DNA required for bacterial biofilm formation. Science (80-.). https://doi.org/10.1126/science.295.5559.1487

Wolff, K.A., de la Peña, A.H., Nguyen, H.T., Pham, T.H., Amzel, L.M., Gabelli, S.B., Nguyen, L., 2015. A Redox Regulatory System Critical for Mycobacterial Survival in Macrophages and Biofilm Development. PLoS Pathog. https://doi.org/10.1371/journal.ppat.1004839

Yamazaki, Y., Danelishvili, L., Wu, M., MacNab, M., Bermudez, L.E., 2006. Mycobacterium avium genes associated with the ability to form a biofilm. Appl. Environ. Microbiol. 72, 819–825. https://doi.org/10.1128/AEM.72.1.819-825.2006

Yang, Y., Thomas, J., Li, Y., Vilchèze, C., Derbyshire, K.M., Jacobs, W.R., Ojha, A.K., 2017. Defining a temporal order of genetic requirements for development of mycobacterial biofilms. Mol. Microbiol. https://doi.org/10.1111/mmi.13734

Zambrano, M.M., Kolter, R., 2005. Mycobacterial biofilms: A greasy way to hold it together. Cell 123, 762–764. https://doi.org/10.1016/j.cell.2005.11.011

