## Supplemental data 1 for "*Mycobacterium abscessus* biofilms produce an ECM and have a distinct mycolic acid profile"

### Slide 1
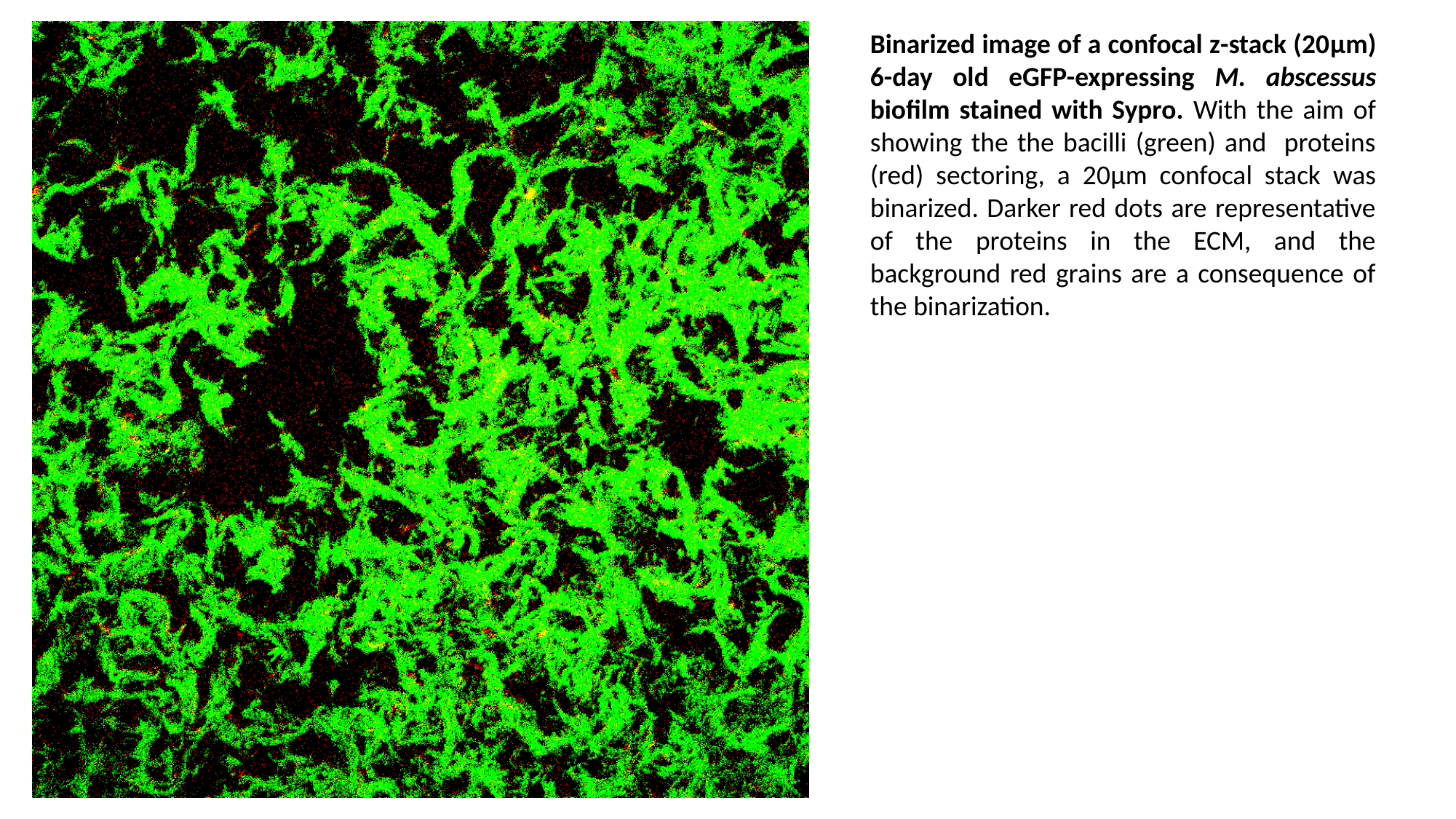

Binarized image of a confocal z-stack (20μm) 6-day old eGFP-expressing M. abscessus biofilm stained with Sypro. With the aim of showing the the bacilli (green) and proteins (red) sectoring, a 20μm confocal stack was binarized. Darker red dots are representative of the proteins in the ECM, and the background red grains are a consequence of the binarization.

### Slide 2
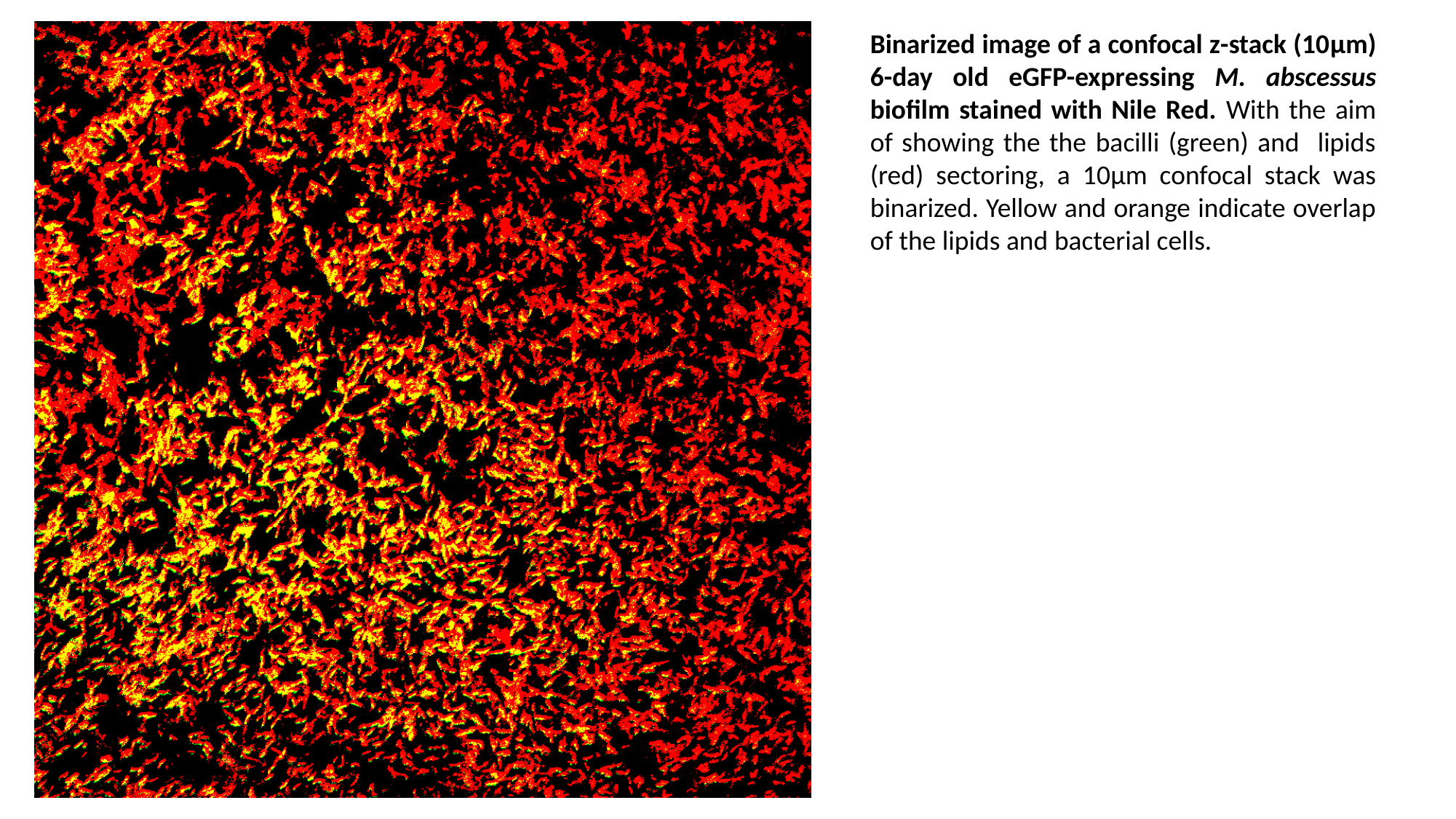

Binarized image of a confocal z-stack (10μm) 6-day old eGFP-expressing M. abscessus biofilm stained with Nile Red. With the aim of showing the the bacilli (green) and lipids (red) sectoring, a 10μm confocal stack was binarized. Yellow and orange indicate overlap of the lipids and bacterial cells.

### Slide 3
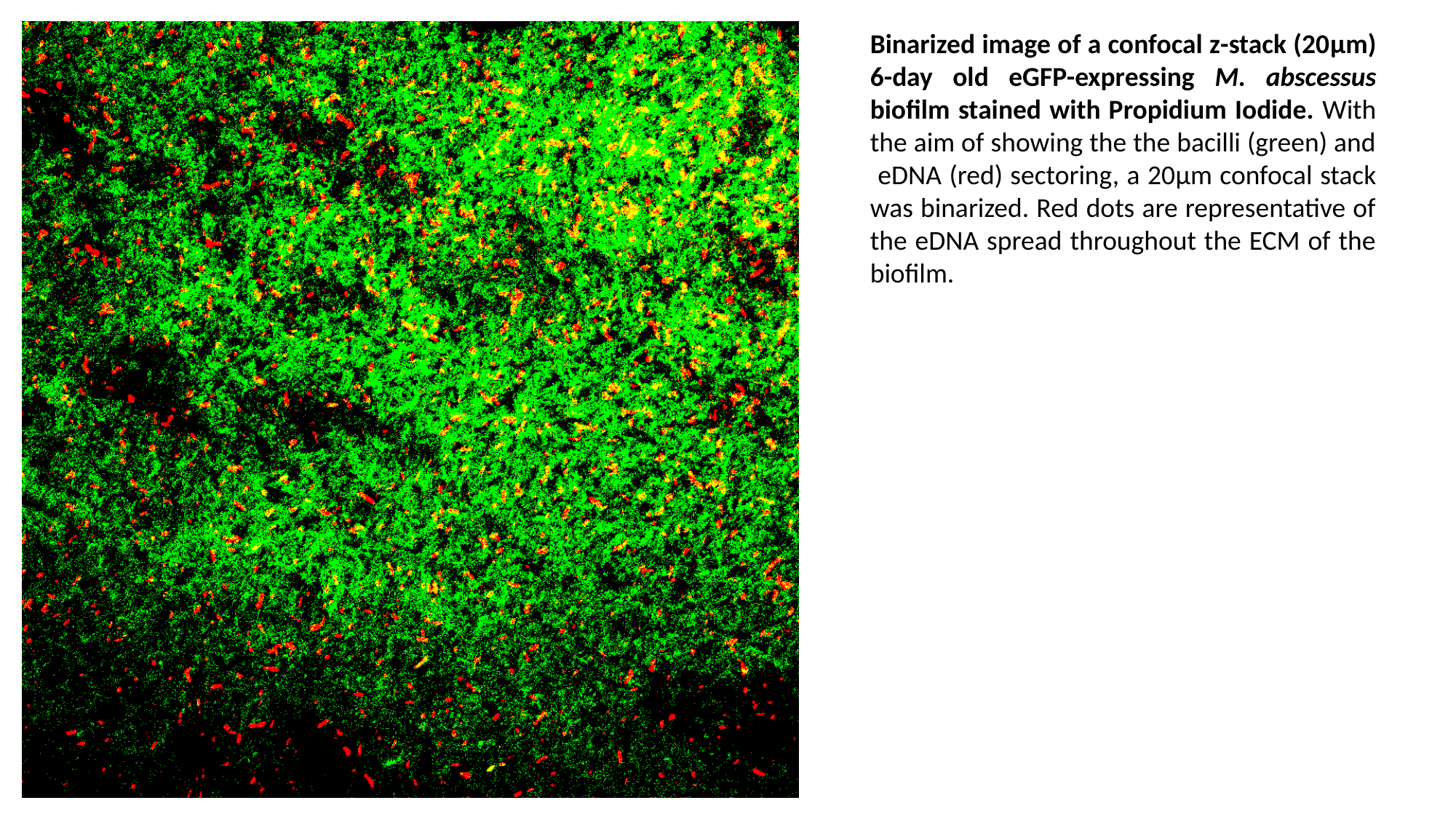

Binarized image of a confocal z-stack (20μm) 6-day old eGFP-expressing M. abscessus biofilm stained with Propidium Iodide. With the aim of showing the the bacilli (green) and eDNA (red) sectoring, a 20μm confocal stack was binarized. Red dots are representative of the eDNA spread throughout the ECM of the biofilm.

### Slide 4
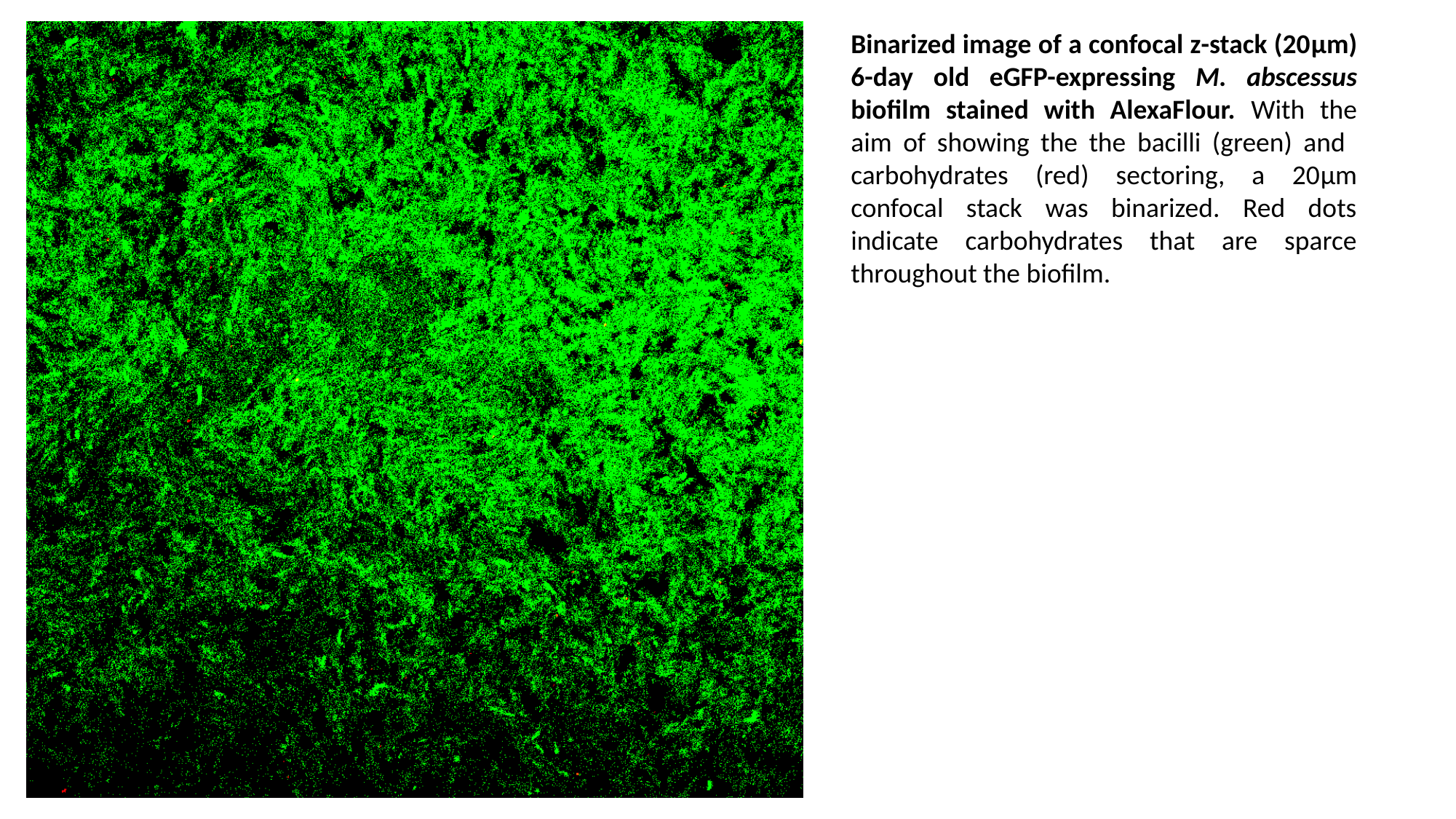

Binarized image of a confocal z-stack (20μm) 6-day old eGFP-expressing M. abscessus biofilm stained with AlexaFlour. With the aim of showing the the bacilli (green) and carbohydrates (red) sectoring, a 20μm confocal stack was binarized. Red dots indicate carbohydrates that are sparce throughout the biofilm.
