## Supplemental data 3 for "*Mycobacterium abscessus* biofilms produce an ECM and have a distinct mycolic acid profile"

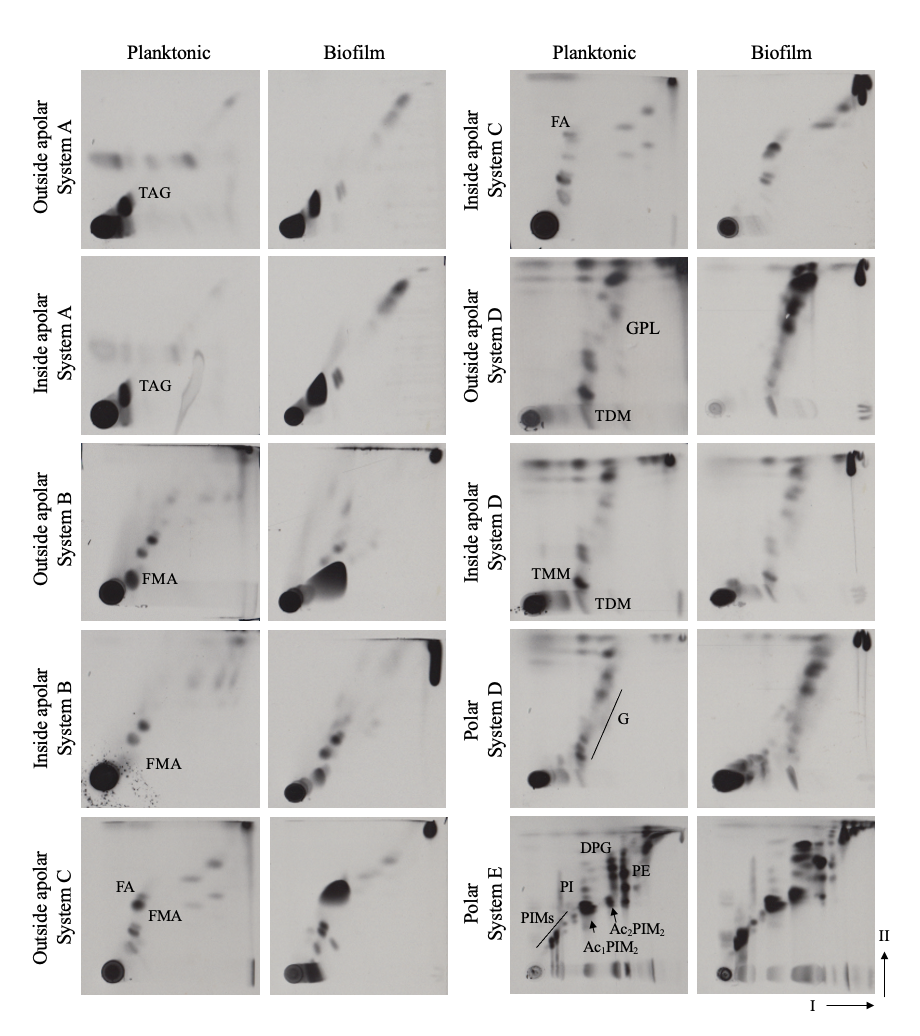
**Supplementary figure 3:** Polar and apolar lipids resolved in various solvent system, listed in Besra, 1998, showing the full lipid profile of *M. abscessus* in planktonic and biofilm cells. TAG, triacylglycerols; FA, fatty acid; FMA, free mycolic acid, TMM, trehalose, monomycolate; TDM, trehalose dimycolate, G,glycolipids; P, phospholipids; DPG, diphosphatidylglycerl; PE, phosphatidylethanolamine; PI, phosphatidylinositol; PIMs, phosphatidylinositol mannosides; Ac_2_PIM_2_, diacyl phosphatidylinositol dimannoside; Ac_2_PIM_6_, diacyl phosphatidylinositol hexamannoside.
