## Supplemental data 4 for "*Mycobacterium abscessus* biofilms produce an ECM and have a distinct mycolic acid profile"

### Slide 1
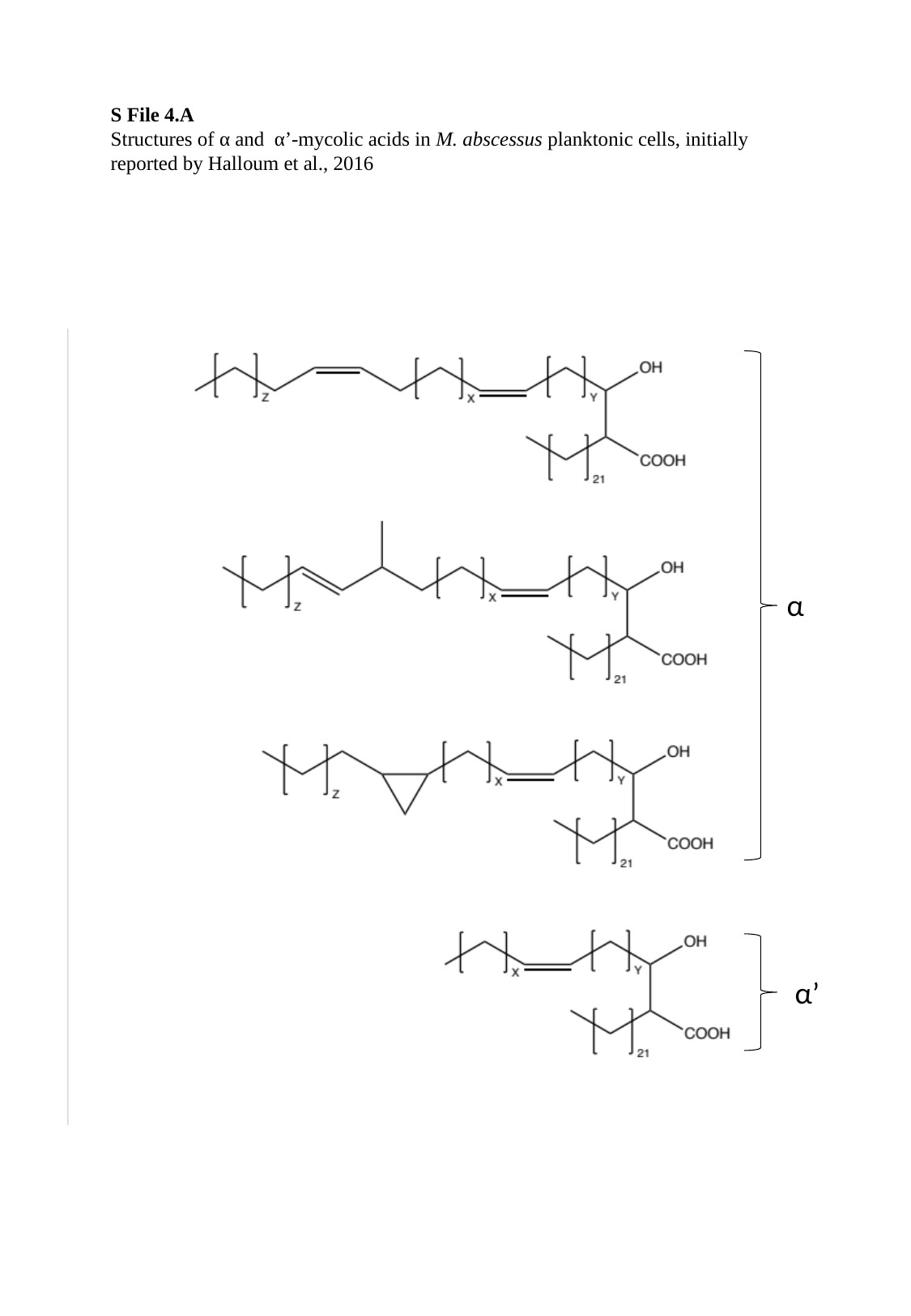

S File 4.A
Structures of α and α’-mycolic acids in M. abscessus planktonic cells, initially reported by Halloum et al., 2016
α
α’

### Slide 2
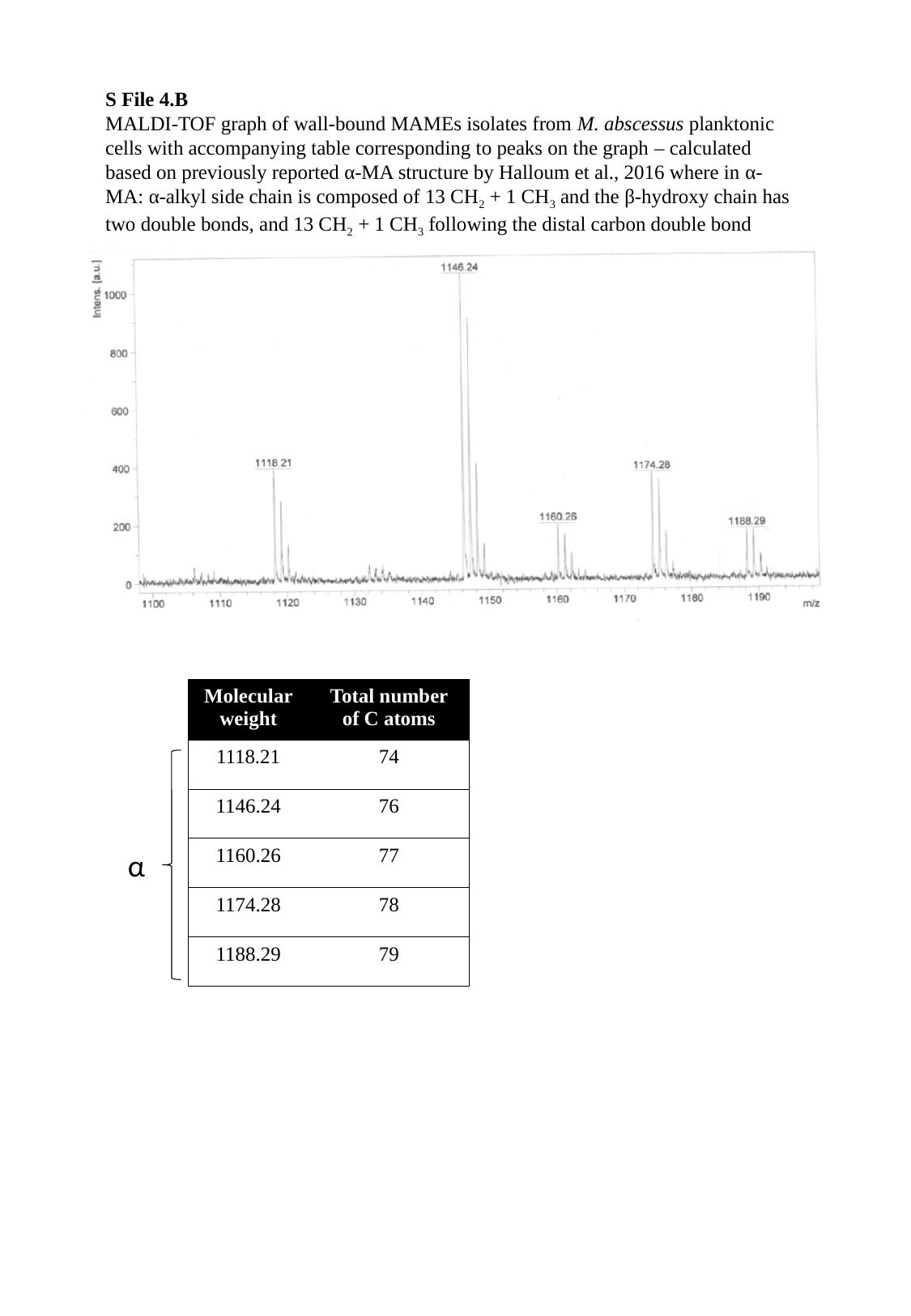

S File 4.B
MALDI-TOF graph of wall-bound MAMEs isolates from M. abscessus planktonic cells with accompanying table corresponding to peaks on the graph – calculated based on previously reported α-MA structure by Halloum et al., 2016 where in α-MA: α-alkyl side chain is composed of 13 CH_2_ + 1 CH_3_ and the β-hydroxy chain has two double bonds, and 13 CH_2_ + 1 CH_3_ following the distal carbon double bond
| Molecular weight | Total number of C atoms |
| --- | --- |
| 1118.21 | 74 |
| 1146.24 | 76 |
| 1160.26 | 77 |
| 1174.28 | 78 |
| 1188.29 | 79 |
α

### Slide 3
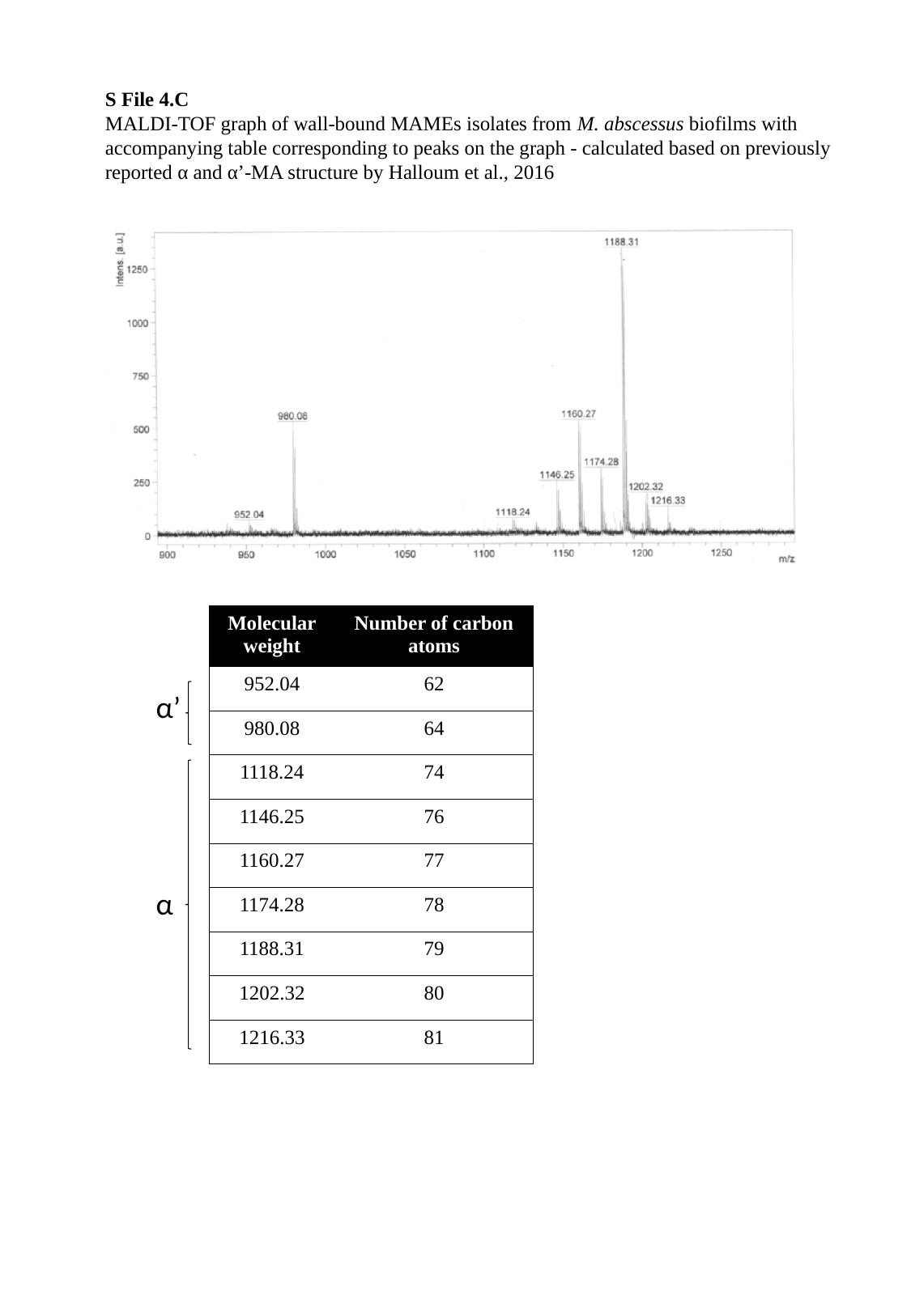

S File 4.C
MALDI-TOF graph of wall-bound MAMEs isolates from M. abscessus biofilms with accompanying table corresponding to peaks on the graph - calculated based on previously reported α and α’-MA structure by Halloum et al., 2016
| Molecular weight | Number of carbon atoms |
| --- | --- |
| 952.04 | 62 |
| 980.08 | 64 |
| 1118.24 | 74 |
| 1146.25 | 76 |
| 1160.27 | 77 |
| 1174.28 | 78 |
| 1188.31 | 79 |
| 1202.32 | 80 |
| 1216.33 | 81 |
α’
α
